# Deepening biomedical research training: Community-Building Wellness Workshops for Post-Baccalaureate Research Education Program (PREP) Trainees

**DOI:** 10.1101/2024.03.10.584300

**Authors:** Dezmond Cole, Andrew S. Eneim, Cory J. White, Chelsy R. Eddings, Morgan Quinn Beckett, Vincent Clark, Jasmin Jeffery, Virangika K. Wimalasena, Alexis Figueroa, Jose Javier Rosado-Franco, Rama Alhariri, Bonita H. Powell, Parris Whitney Washington, Lana N. Christensen, Katherine L. Wilson, Jennifer M. Kavran

## Abstract

**Problem:** All trainees, especially those from historically minoritized backgrounds, experience stresses that may reduce their continuation in science, technology, engineering, math, and medicine (STEMM) careers. The Johns Hopkins University School of Medicine is one of ∼45 institutions with a National Institutes of Health funded Postbaccalaureate Research Education Program (PREP) that provides mentoring and a year of fulltime research to prepare students from historically excluded groups for graduate school. Having experienced the conflation of stresses during the COVID-19 pandemic and related shutdown, we realized our program lacked a component that explicitly helped PREP Scholars recognize and cope with non-academic stresses (financial, familial, social, mental) that might threaten their confidence and success as scientists and future in STEMM.

**Intervention:** We developed an early-intervention program to help Scholars develop life-long skills to become successful and resilient scientists. We developed a year-long series comprised of 9 workshops focused on community, introspection, financial fitness, emotional intelligence, mental health, and soft-skills. We recruited and compensated a cohort of PhD students and postdoctoral fellows to serve as Peer Mentors, to provide a community and the safest ‘space’ for Scholars to discuss personal concerns. Peer Mentors were responsible for developing and facilitating these Community-Building Wellness Workshops (CBWW).

**Context:** CBWW were created and exectued as part of the larger PREP program. Workshops included a PowerPoint presentation by Peer Mentors that featured several case studies that prompted discussion and provided time for small-group discussions between Scholars and Peer Mentors. We also included pre- and post-work for each workshop. These touch-points helped Scholars cultivate the habit of introspection.

**Impact:** The CBWW exceeded our goals. Both Peer Mentors and Scholars experienced strong mutual support, and Scholars developed life-long skills. Notably, several Scholars who had been experiencing financial, mental or mentor-related stress immediately brought this to the attention of program leadership, allowing early and successful intervention. At the completion of CBWW, PREP Scholars reported implementing many workshop skills into practice, were reshaping their criteria for choosing future mentors, and evaluating career decisions. Strikingly, Peer Mentors found they also benefitted from the program as well, suggesting a potential larger scope for the role of CBWW in academia.

**Lessons Learned:** Peer Mentors were essential in creating a safe supportive environment that facilitated discussions, self-reflection, and self-care. Providing fair compensation to Peer Mentors for their professional mentoring and teaching contributions was essential and contributed meaningfully to the positive energy and impact of this program.

## INTRODUCTION

The recruitment and retention of individuals from communities that are underrepresented (UR) (including low-income, first-generation college, or rural) due to historical exclusion from careers in science, technology, engineering, math, and medicine (STEMM) is recognized as a national priority for our workforce to remain competitive and to reduce gross disparities in health and healthcare^1–3^. Achieving this goal, however, is challenged by many factors including the long-standing economic, educational and personal impacts of systemic and institutional racism in the USA^4^.

Furthermore, bias and discrimination, which has historically been ingrained within academic institutions, contributes to declines in representation of UR individuals in pursuing academic roles^5^. College-level interventions, especially programs encouraging participation in STEMM research, have increased the retention of UR individuals in academia^6^. Within this landscape, the National Institutes of Health (NIH) funds 45 Postbaccalaureate Research Education Programs (PREPs) nationwide for UR college graduates interested in PhD careers with the goal of increasing entry into PhD programs. PREP programs typically fund 4-6 participants for a year of fulltime research experience, mentoring and professional development, and have proven highly successful in preparing trainees for the scientific and academic demands of graduate programs and PhD careers. Indeed, PREP programs at six institutions in the mid-Atlantic region report a collective success rate of 85% for entry into PhD or MD/PhD programs, and a total of 93 PhDs earned as of 2022^7^.

However, success does not always come easily. PREP Scholars often experience financial stress, mental stress, stereotype-threat or imposter syndrome, any of which can sap confidence and jeopardize continuation in science. Our program previously relied on the program director or scientific mentors to notice stress, or Scholars to bring it up. This approach was inadequate because these kinds of stress can be internalized by high-achieving students, remaining ‘invisible’, unrecognized or unacknowledged.

The experiences of our PREP Scholars mirror national reports that graduate students from historically UR groups are more likely than their white peers to report insufficient financial resources and to have higher rates of mental health struggles such as anxiety, depression and suicidal ideation, likely reflecting the higher accumulated burden of systemic racism, sexism, harassment and micro-aggressions faced by these students^4,8–12^. Burnout, anxiety, depression, work-life imbalance, social isolation and imposterism are exacerbated by habits of overwork and overachievement by UR individuals in response to systemic discrimination in our society^13^.

To address these issues head-on, we needed to help Scholars build emotional skills and support-systems essential for personal resilience and long-term success in STEMM research. We designed a series of Community-Building Wellness Workshops (CBWW) that included a peer mentoring system to help Scholars (a) identify potential stressors, (b) develop positive coping skills to prevent or address challenges as they arise, (c) normalize asking for help, and (d) develop a community support system with a cohort of PhD students and postdoctoral fellows that functions as Peer Mentors (Peer Mentors). By incorporating these monthly workshops into the existing PREP curriculum, we made wellness a new core component of our program. Our first cohort of PREP Scholars participating in CBWW reported implementing these skills in their daily lives, and the majority said this training strengthened their criteria for choosing compatible mentors and evaluating career paths. We report here our experiences, lessons learned, and provide ADA-accessible slides and worksheets for all nine workshops for use by other programs (***Supplemental Information S1-S13***).

## PROGRAM DESIGN

### Context of the Program

We designed CBWW to mesh with our existing NIH-funded PREP at Johns Hopkins. Hopkins PREP provides full-time paid research experience and mentoring to aspiring scientists from backgrounds historically excluded in science who are aiming for entry and success in competitive PhD programs. Each year we accept a cohort of ∼12 recent college graduates each year for a planned one-year (if aiming for PhD) or two-year (if aiming for MD/PhD) program that combines discovery-based research in a university lab with custom mentoring and skill-building. Recognizing that our Scholars had unmet needs exacerbated by the COVID-19 pandemic, we obtained an NIH supplement to expand the scope of our program to holistically nurture Scholars by addressing personal and emotional well-being and teaching that these life skills are essential to thrive and persist in academia.

We had two major goals that went hand-in-hand: to create a community with more-experienced peers and, within this safe community, provide wellness skills in five core areas: introspection, financial fitness, mental health, emotional intelligence, soft skills, introspection (***Figure 1, Table 1***). In addition to equipping Scholars with skills needed by resilient scientists, we hoped these workshops would help Scholars to identify any ongoing stress they were experiencing and to normalize seeking help thereby facilitating early intervention. Introspection was a core principle of each workshop, with case studies and questions to guide reflection on various aspects of their personal and professional lives. Peer Mentors were pivotal in this endeavor, providing community and camaraderie while serving as role models. They shared their own journeys through academia that allowed them to empathize and relate to Scholars and create a safe space for confidential dialogue and discussion.

**Table 1.** Workshop Topics and Themes.

| **Workshop** | **Order in Series** | **Theme** | **Created and Facilitated by** |
| --- | --- | --- | --- |
| Values & Goals | 1 | Introspection | ASE |
| Owning your Identity | 9 | Introspection | JJ and CJW |
| Mental Health Strategies | 8 | Mental health | DC and JJR-F |
| Identifying Stressors | 3 | Mental health | MQB and PWW |
| Imposter Syndrome & Racial Bias | 4 | Mental health | CJW |
| Financial Skills | 2 | Financial Fitness | CRE |
| Difficult Conversations | 6 | Emotional intelligence | LNC and VKW |
| Self-Advocacy | 7 | Emotional intelligence | RA and AF |
| Time management | 5 | Soft skills | VC and BHP |
| All of the above | n/a | Community | RA, MQB, DC, LNC, VC, C.R.E., A.S.E., AF, JJ, JMK, BH, JJR-F, PWW, CJW |

**Figure 1.**
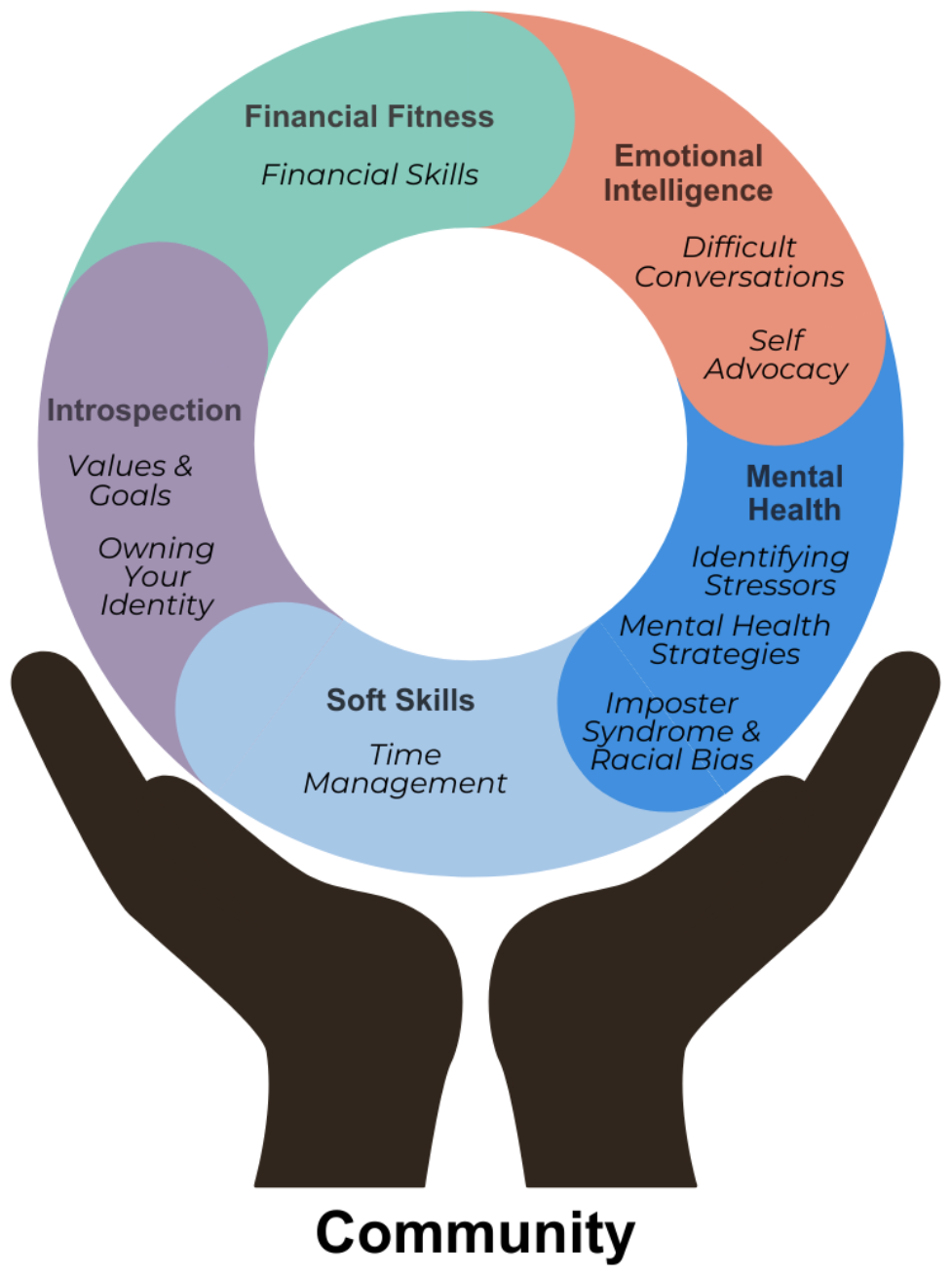
Holis&c Wellness Training for Scholars. An illustra+on of the concept and design for the CBWW. Individual workshops are clustered under five themes: Financial Fitness, Emo+onal Intelligence, Mental Health, SoE Skills, and Introspec+on with individual workshops indicated in italics.

### Specific Interventions

#### Peer Mentors

To create a broader mentoring community for PREP Scholars than previous, which previously had included the PREP leadership, a “mini-thesis” committee of 3 faculty, and their lab mentor and labmates, we choose to recruit a team of Peer Mentors who would help develop CBWW and serve as the foundation of the program. We solicited applications from the entire population of PhD and MD/PhD students and postdoctoral fellows at the School of Medicine and received 38 applications for 14 positions. This widespread interest allowed the workshop organizer. PREP associate director (JMK), and PREP director (KLW) to select a broadly diverse group of Peer Mentors committed to increasing diversity in science. We recruited 14 inaugural Peer Mentors, including several with parallel journeys (alumni from PREP or equivalent programs), all with previous experience in mentoring, and many from historically excluded backgrounds. Peer Mentors, all of whom co-authored this report, were fundamental to the creation and success of this wellness program. The Peer Mentors convened together at the start of the year with CBWW organizer (JMK) to develop a list of workshop topics and decide who would be responsible for developing and facilitating each. They poured their experiences, academic journeys, insights and passion into selecting topics, then worked to create and personally facilitate ‘their’ workshop, usually in pairs, with messages that would resonate with Scholars, while forging bonds of understanding and camaraderie. Recognizing that sustained peer-mentoring deserves to be compensated as a professional activity, we paid Peer Mentors as teaching assistants through funds provided by an NIH^10,14^.

Each month, one or two Peer Mentors facilitated the workshop they designed, while other Peer Mentors participated in the audience and led small group discussions with Scholars (***Table 1***). Outside these sessions, Peer Mentors soon began engaging informally with PREP Scholars, for example, meeting for coffee or lunch, for further guidance and support. Importantly, many potential problems were brought up during these informal sessions. In consultation with their Peer Mentors, several Scholars developed action plans that included bringing their issue to the attention of their research mentor or PREP program leadership.

#### Wellness Workshops

Workshops were held monthly and followed a similar format designed to build community, trust, and cultivate a habit of wellness. Each 60-minute workshop included ample time (up to 30 minutes) for small-group discussions, usually involving 4-6 PREP Scholars and 2-3 Peer Mentors. Peer Mentors guided these discussions, drawing on their own journeys and knowledge about institutional and other resources, and ensuring that discussions remained relevant and resonated with the Scholars’ lived experiences. Small groups allowed intimate discussions, allowing every voice to be heard. Additionally, there was protected time after each workshop to hang-out in the same room and allow for small group conversations to continue and as an additional opportunity for Scholars to interact with each other and Peer Mentors. To normalize the practice of wellness, each workshop encouraged Scholars to introspect, and provided opportunities for those who were comfortable doing so to share their insights. We also assigned two short exercises, communicated via email from the workshop organizer, to be completed either two weeks before or after each workshop. These “touch points” (included in the slides for each workshop, ***Supplemental Information S1-S9, S12-13***) reinforced key points from the workshops and routinely setting aside time for self-work.

We felt that the success of this program depended on building trust so that allow all participants felt comfortable sharing their experiences and becoming part of the community. The first workshop included a discussion about trust and active listening.

Throughout the year we often referred to this discussion, especially when starting a new workshop that might be triggering, and whenever anyone shared a particularly vulnerable experience with their group. To preserve the sanctity of the Peer Mentor/Scholar dynamic, we decided not to invite external speakers to serve as workshop facilitators; instead, Peer Mentors facilitated the same workshop they had created and only one faculty member was present (workshop organizer JMK).

We ran one workshop each month during the academic year that covered nine topics associated with our overarching themes of introspection, financial fitness, mental health, emotional intelligence, and soft skills (***Table 1, Figure 1*)**. Specific topics were chosen in response to past experiences of Peer Mentors and PREP program leaders; we endeavored to address many of the multifaceted challenges Scholars may face when navigating academia and life beyond, but do not claim to have addressed all important topics. The resources developed for each workshop are provided in ***Supplemental Information (S1-S13)***, with our main themes described and summarized below.

To promote **Introspection**, we introduced Scholars to the practice of self-reflection during the kick-off workshop, “*Goals and Values*”, and reinforced this practice in the final workshop, “*Owning Your Identity*”. During “*Goals & Values*”, Scholars were tasked with evaluating whether or how their professional goals align with their personal values. This same lens was used to navigate the rest of the program and was referred to frequently in other workshops. The “*Owning Your Identity*” workshop helped Scholars evaluate how their professional responsibilities, family obligations, financial obligations and social life all intertwine with well-being, so they could adopt a growth mindset as these factors change over time. This workshop also encouraged Scholars to reflect on their scientific progress, self-acknowledge real achievements, and think ahead about their future professional endeavors. Each Scholar also created and shared a slide representing their values with the group to help build community; these slides were so revealing and fun that ‘group shares’ were repeated in several other workshops.

**Financial Skills**, covered minimally in one workshop, included basic elements of budgeting, strategies for saving, key points about student loans, and fee waivers for graduate school applications. This workshop mainly focused on identifying and solving any current financial stress, and developing positive skills and strategies to ensure Scholars can live comfortably within their income and start building long-term financial stability. This workshop was particularly useful for Scholars who carry high student loan debt; on average, 49% of PREP Scholars over the last 5 years carry student loan debt.

Three workshops supported **Mental Health**: “*Mental Health Strategies*”, “*Identifying Stressors*”, and “*Intersection of Imposter Syndrome and Racial Bias*”. Our overarching goals for these workshops were to destigmatize mental health issues by providing tools to help Scholars manage stress and anxiety, develop resilience, and normalize the process of seeking professional mental help. Many descriptions of imposter syndrome neglect the impacts of racial and gender bias as major contributors to imposterism; this leaves researchers from backgrounds historically excluded in science at elevated risk of self-doubt, stereotype threat, perceived incompetence, or sense of non-belonging^15,16^. We therefore developed a workshop that explicitly explored this linkage, drawing heavily on Peer Mentor experiences, for our workshop on “*Imposter Syndrome and Racial Bias*.” This workshop, the fifth in our series, when scholar participation erupted, and everyone began talking. To equip Scholars with a strategy to work through their own solutions, at the end of this workshop we introduced Peer Coaching Groups. This activity was adapted from the University of Washington’s

Broadening the Representation of Academic Investigators in NeuroScience program organized by Dr. Claire Horner-Devine, Dr. Cara Margherio, Dr. Sheri Mizumori and Dr. Joyce Yen^17^. The activity consisted of concretely identifying your dilemma, being questioned by a mentor or another scholar, and then forming an actionable plan. The goal of this exercise was to equip Scholars with another tool to self-reflect, identify sources that may contribute to imposterism, and strategize ways to address them. “*Identifying Stressors*” took a practical approach to understanding how stress manifests both emotionally and physically, and included coping strategies to relieve stress. “*Mental Health Strategies*” normalized the practice of self-care and seeking mental health treatment as essential in demanding academic environments.

Two **Emotional Intelligence** workshops helped Scholars develop interpersonal skills: “*Difficult Conversations*” helped Scholars learn how to evaluate different kinds of conflict and develop a plan to resolve conflict, with the goal of maintaining healthy and effective working relationships. After defining what a difficult conversation in biomedical research may look like, the Scholars were given a framework based on worksheets from the Johns Hopkins Ombuds for Doctoral and Postdoctoral Students and Fellows Program, Annalisa Peterson, to evaluate how to proceed with having a courageous conversation if necessary. This framework included identifying an appropriate person to talk to, initiating the conversation, using “I” statements to strategically have the conversation, being open to discussion and new information, and identifying solutions and confirming accountability measures. In “*Self-Advocacy*”, Scholars examined different communication styles, and practiced using assertive communication to define boundaries. Scholars also worked on their emotional intelligence through case studies that illustrated how to improve communication, develop stronger relationships, and resolve conflicts. Teaching self-advocacy skills helps minoritized students succeed academically and gain confidence in their own identity and their place in science, thereby increasing their likelihood of academic success^18,19^.

For **Soft Skills**, we developed one workshop on **“***Time-management*”. The pre-work for this workshop asked Scholars to critically evaluate their current time-management skills, and then provided concrete strategies to manage time (such as Gantt charts, weekly planners, and the Pomodoro Technique)^20,21^. To align these practices with their personal goals, the workshop included examples relevant to the common needs of Scholars (i.e. creating a three-month schedule, an action plan for an experiment, or an MCAT study timeline). Small-group discussions emphasized self-care (***not*** self-sacrifice) as a fundamental part of time-management and, to link back to previous themes, explored how stress may manifest when dealing with deadlines and talking about practices of self-care to help reduce that stress. This workshop exemplifies how we tried to reinforce multiple themes by linking to other workshops.

### IMPACT

At the end of the year, we evaluated the efficacy of the CBWW by surveying PREP Scholars and Peer Mentors (***Figures 2, 3, S14-S17***). Our response rate was 86% of Scholars, 100% of Peer Mentors. Although it is still too early to gauge long-term impact, our short-term results were highly encouraging and warrant continuing these Workshops.

**Figure 2.**
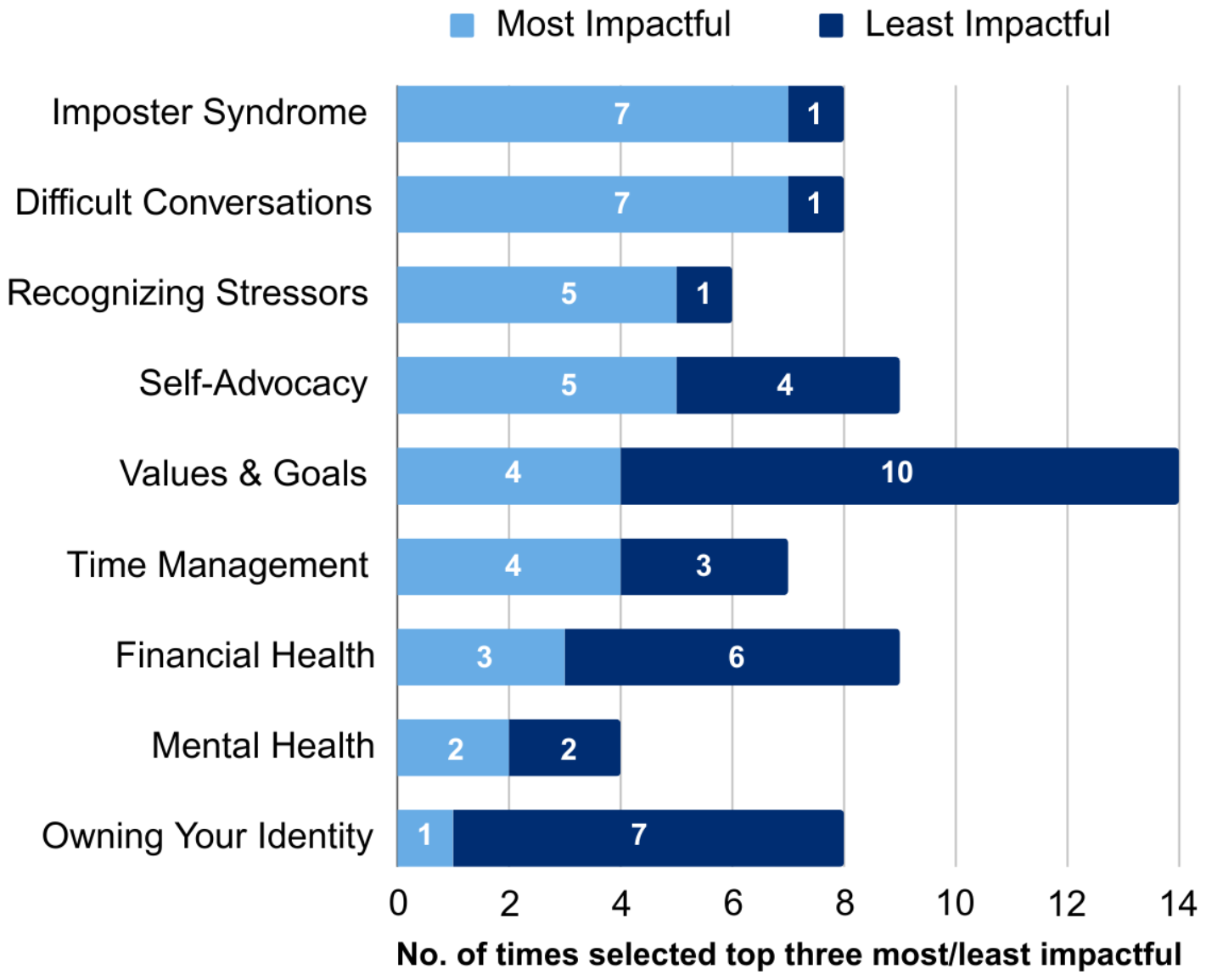
Workshop impact. In the exit survey, Scholars selected the three most and least impacHul workshops. The chart shows the number of +mes a workshop was selected as each with the most impacHul indicated in light blue and least impacHul in dark blue.

**Figure 3.**
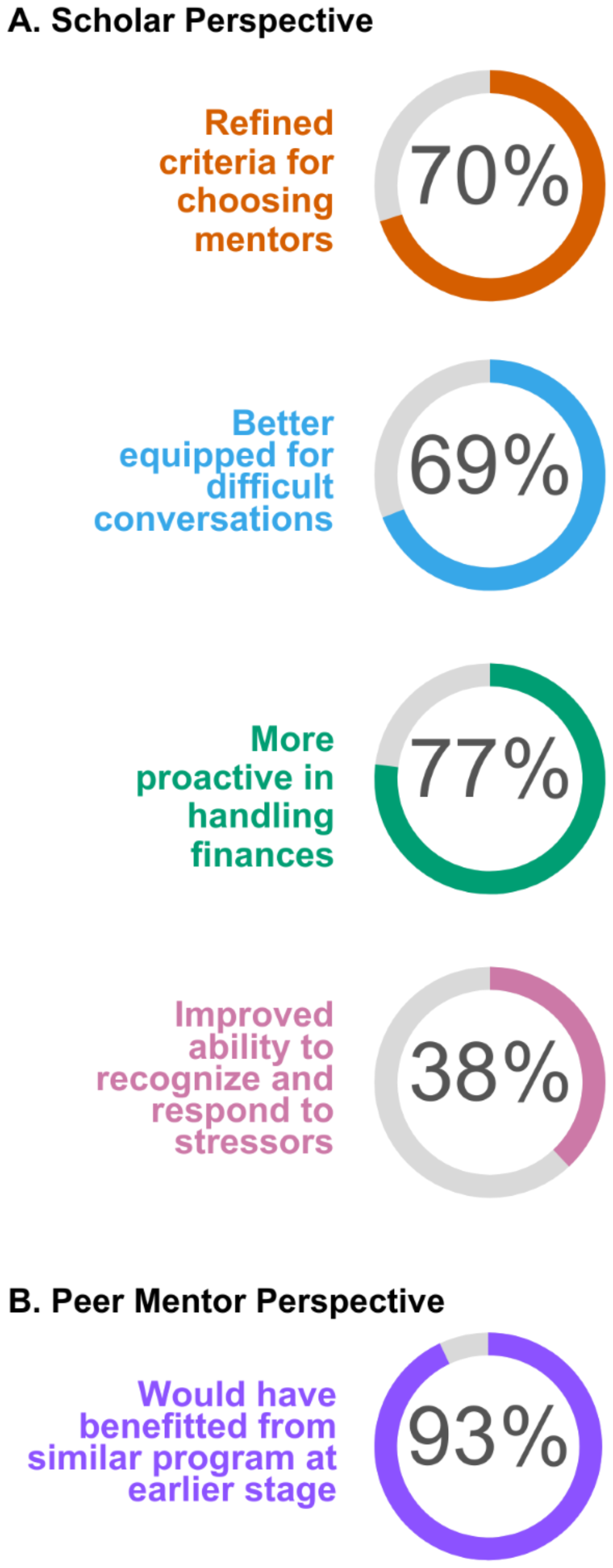
Impact of Wellness Program. Donut charts based on survey results highligh+ng key impacts of the CBWW on for the Scholars (A) and Peer Mentors (B).

When asked to identify their three most impactful workshops, we had multiple tied scores (***Figure 2***). Scholars had two clear favorites: *Imposter Syndrome* and *Difficult Conversations*, each named by 7 of 13 Scholars (53.8% each), closely followed by *Recognizing Stressors* and *Self-Advocacy* (38.5% each), followed by *Values and Goals* and *Time Management* (31%). The three least impactful workshops were *Values and Goals* (76.9%), *Owning your Identity* (53%), and *Financial Health* (46%).

Interestingly, three workshops had opposite impacts, being highly valued by some Scholars and unvalued by others (*Self-Advocacy, Values & Goals, Time Management*). The varied impacts of the same workshop was unsurprising given the unique combinations of skillsets and needs among PREP Scholars. We infer that the *Mental Health* workshop was appreciated by the majority of Scholars, even though it did not generate strong feelings: it was deemed most-impactful by only two Scholars and least-impactful by only two other Scholars. This discrepancy may reflect that several Scholars already shared that they already had good mental hygiene including seeing a therapist which could lessen the impact of these workshops for those Scholars. When considering the core values embodied by each workshop, we realized that overall the most impactful workshops concerned mental health and emotional intelligence (e.g., developing strategies to navigate scientific spaces and professional relationships), whereas the least impactful both focused on introspection. These introspection workshops, *Values and Goals* and *Owning your Identity*, were the most conceptual and did not provide the ‘instant’ tangible skills offered by other workshops which may explain their lower impact.

#### Impact on Scholars

During the first few workshops, a common refrain from Scholars was that they had never really thought about these topics before, suggesting they were unfamiliar with wellness or introspection. Nearly half of PREP Scholars are first-generation students, and first-generation students frequently deal with higher levels of mental health challenges reflecting a lack of familial support to help navigate these spaces^22,23^. At the end of the year, however, 85% of Scholars reported putting workshop topics into practice, suggesting a tangible impact on their professional lives (***Supplemental Figure S16)***. Just under half of our Scholars reported changing how they interact with their advisors and lab mates because of the workshops (***Supplemental Figure S16***). The majority of Scholars reported feeling better prepared for multiple aspects of their future careers: 69% felt better equipped to have difficult conversations in a professional setting; 38% reported an improved ability to recognize when a situation is becoming stressful enough to take action, and 77% felt more proactive in handling their finances (***Figure 3A***).

Other survey results suggest Scholars acquired skills that will serve them well as scientists. 70% of Scholars reported that they refined their criteria for choosing future mentors, a potentially long-term impact on success in graduate school and beyond (***Figure 3B)***. Based on our “*Difficult Conversations*” Workshop, Scholars reported feeling significantly better equipped to raise concerns with faculty, lab members, and peers, and 85% reported a better understanding of assertiveness after our workshop on self-advocacy that we interpret as a clear sign they had begun building skills as professionals to cope with the stresses encountered in STEMM settings (***Supplemental Figure S16***).

Scholars responded overwhelmingly that their willingness to share during workshops was due to the openness of the Peer Mentors, who shared their own lived experiences. By the end of the year, 38% reported feeling completely comfortable sharing personal experiences during workshops with 46% feeling very comfortable and 23% feeling moderately comfortable (***Supplemental Figure S17***). We conclude that Peer Mentors were both essential and successful in building a community where Scholars felt supported both professionally and personally. We are hopeful that Scholars will amplify this training as future Peer Mentors to help others.

#### Impact on Peer Mentors

Peer Mentors also reported benefiting from the CBWW. Peer Mentors were nearly unanimous (93%) in reporting that they would have personally benefited from similar programming at this stage or earlier in their training (***Figure 3B***). Therefore, one important take-home message at the institutional level (particularly PhD training programs) is that wellness workshops doubled the intended impact by benefitting the trainees who served as Peer Mentors at least as much as they benefitted PREP Scholars. Since we did not anticipate this strong dual impact, better assessment tools will be needed in future to evaluate and understand how these workshops impact Peer Mentors.

#### Impact on the PREP program

We frequently observed that different sets of PREP Scholars, typically 4-6, stayed and sought out Peer Mentors to discuss an ongoing situation informally, after the workshop ended. Since these discussions were confidential, the Peer Mentors provided critical insights and feedback; in at least four cases, this included urging the scholar to take the next step towards early intervention. Indeed, 25% of Scholars reported having discussed topics covered in the workshops with either their lab mentor or the PREP program director; those who approached the PREP director discussed an issue raised in a specific workshop that was directly affecting their training, well before they had reached a crisis point, allowing timely and appropriate intervention (***Supplemental Figure 16)***. These events each took place within five days after the relevant workshop. From the point of view of Peer Mentors and program leadership, this truly demonstrated that Scholars had learned to identify potential stressors, tackle difficult conversations and self-advocate.

Overall, this programming was intended to facilitate habits of introspection and provide tangible tools that Scholars may not have developed in a systematic way without these workshops. Ultimately, these workshops were effective in helping Scholars feel better equipped to enter the next phase of their career: 100% of Scholars who participated felt they benefitted from these workshops (***Supplemental Figure S16***).

### LESSONS LEARNED

Our final analysis of these workshops surprised us: the specific topics covered were less important; instead, the most important thing was that these workshops created a safe, confidential, and supportive space for Scholar-led discussion. The Peer Mentors were essential to create and maintain this professionally and personally safe and supportive community for the Scholars. Although some workshops certainly ‘resonated’ with Scholars, they had no clear overall preference in terms of ‘most vs least’ impactful workshops, but instead stressed that the best and most valuable components of the program were their interactions with Peer Mentors. Scholar feedback revealed that the willingness of Peer Mentors to be vulnerable about their own experiences was by far the most helpful aspect of the program for the Scholars. Peer Mentors brought something infinitely precious to these workshops: their own lived experiences and dedication to help others walk their pathway with confidence and joy.

By normalizing their shared experiences, Peer Mentors helped Scholars learn how to navigate spaces in STEMM that are not sufficiently diverse. When Peer Mentors shared their struggles, Scholars realized they are not alone if they feel overwhelmed. With Peer Mentors responsible for leading these workshops and discussions, Scholars were able to visualize a future in which they too can succeed in STEMM. To forge this bond, it is critical for Peer Mentors to directly reflect the racial, ethnic, socioeconomic and cultural diversity of the Scholars.

Small group discussions based on case-studies were a core component of the shared format of these workshops because they fostered discussion and introspection. We observed that Scholars were most engaged when case studies were used as starting points for small group discussions. Case studies often spurred Scholars to apply the hypothetical situation to their own experiences, and work through their responses with their small group. This format also allowed Peer Mentors to flexibly adapt the topic to the current needs of the Scholars in their small group. Peer Mentors used these opportunities to urge Scholars to think more deeply about the situation and consider different ways they could handle it. Scholars agreed these discussions were a highlight of the program and requested more opportunities to interact with Peer Mentors beyond these formal workshops, to which we responded with scheduled, optional coffee-hours or lunch-dates at a nearby café.

To implement this type of programming at other institutions, we suggest focusing first on identifying potential Peer Mentors. Funding to cover their professional time at the level of Teaching Assistants is also essential for recruitment. Peer Mentors should be near peers, ideally representing the next two steps appropriate for STEMM progression in their field (e.g., Masters students, PhD students and/or postdoctoral fellows), who have shared experiences with the Scholars. Two additional aspects to consider are that Scholars should have multiple and regular opportunities to interact with Peer Mentors, both formally and especially informally, and that workshops should include significant time for small group discussions, which gradually builds increasing trust within the community that is most likely to translate into deeper and more impactful discussions during workshops. Our experiences suggest CBWW would build and strengthen community and benefit trainees at each step of their career (including undergraduates, Masters, and PhD students). Our need for early intervention with Scholars lends evidence that this type of programming at increasingly earlier stages of training could provide dividends and be a critical contributor to future success. In this first year we had nearly a 1:1 ratio of PM to Scholars (14:16). This large cohort of Peer Mentors allowed for occasional absences by some at any given workshop but still allowed for 1-2 Peer Mentors to lead each small group discussion. We could not find a specific study on the optimal mentor to mentee ratio, but generally more mentors are found to be better^24,25^.

We are continuing CBWW as an integral part of the PREP program and strongly encourage other institutions to embrace and adapt these principles into their training. Early feedback and experience demonstrate that these workshops were successful in preventing early challenges from derailing PREP Scholars’ futures in the short term and in equipping Scholars with skills essential for resilience as scientists in the long term.

We feel that programming of this nature can play a pivotal role in addressing the persistent challenges faced by historically underrepresented groups in STEMM.

## Supporting information

S2 Slidedeck for Financial Values Workshop

S3 Slidedeck for Identifying Stressors Workshop

S4 Slidedeck for Mental Health Strategies Workshop

S7 Slidedeck for Self Advocacy Workshop

S8 Slidedeck for Racial Bias and Imposter Syndrome

S9 Slidedeck for Difficult Conversations

S12 Slidedeck for Time Management

S13 Slidedeck for Owning your Identity

S14 Scholar Exit Survey Questions

S15 Peer Mentor Exit Survey Questions

S16 Survey Results from Scholars

S17 Survey Results from Peer Mentors

S1 Slidedeck for Values & Goals Workshop

## ACKNOWLEDGEMENTS

We would like to thank Annalisa Peterson, Ombuds for Doctoral and Postdoctoral Students, Fellows, and Programs at Johns Hopkins University, for providing the worksheets on Difficult Conversations.

## SUPLEMENTARY MATERIALS

S1 Slidedeck for Values & Goals Workshop S2 Slidedeck for Financial Values Workshop

S3 Slidedeck for Identifying Stressors Workshop

S4 Slidedeck for Mental Health Strategies Workshop

S5 Worksheet for Mental Health Strategies – Self Care Checkup

S6 Worksheet for Mental Health Strategies – Daily Mood Tracker

S7 Slidedeck for Self Advocacy Workshop

S8 Slidedeck for Racial Bias and Imposter Syndrome

S9 Slidedeck for Difficult Conversations

S10 Worksheet for Difficult Conversations – Courageous Conversations Discussion Planner

S11 Worksheet for Difficult Conversations – Courageous Conversations Reflections

S12 Slidedeck for Time Management

S13 Slidedeck for Owning your Identity

S14 Scholar Exit Survey Questions

S15 Peer Mentor Exit Survey Questions

S16 Survey Results from Scholars

S17 Survey Results from Peer Mentors

