## Supplementary material for "Deepening biomedical research training: Community-Building Wellness Workshops for Post-Baccalaureate Research Education Program (PREP) Trainees": S2 Slidedeck for Financial Values Workshop

### Overview

- The goal of this workshop is to increase Scholar long-term financial literacy by providing concrete resources and discussion from which they can start/continue their own research into their individual financial situation (slides should be shared as they contain links)

- We do not want to tell them how to spend their money; we want them to reflect on how they currently handle their money and if they will do anything different after this discussion

### Pre-work: speed budgeting exercise

**Building a basic budget relies on 3 things:**

1. Knowing your net income (from all sources)
2. Creating a list of your monthly expenses
3. Calculating a monthly amount of income left over (your “cash flow”)

https://www.huntington.com/learn/budgeting/how-to-make-a-budget

**Use the provided worksheet and fill out the following info in ONLY 3 minutes, then be prepared to discuss the day of the workshop:**

- Create a list of your monthly expenses (specific $ amount NOT needed), identify which are fixed vs. variable
  - Fixed: same amount each month
  - Variable: amount can fluctuate each month
- Characterize which are “want” vs. “need” expenses (waying your priorities)
  - Want: trip to Bali
  - Need: groceries

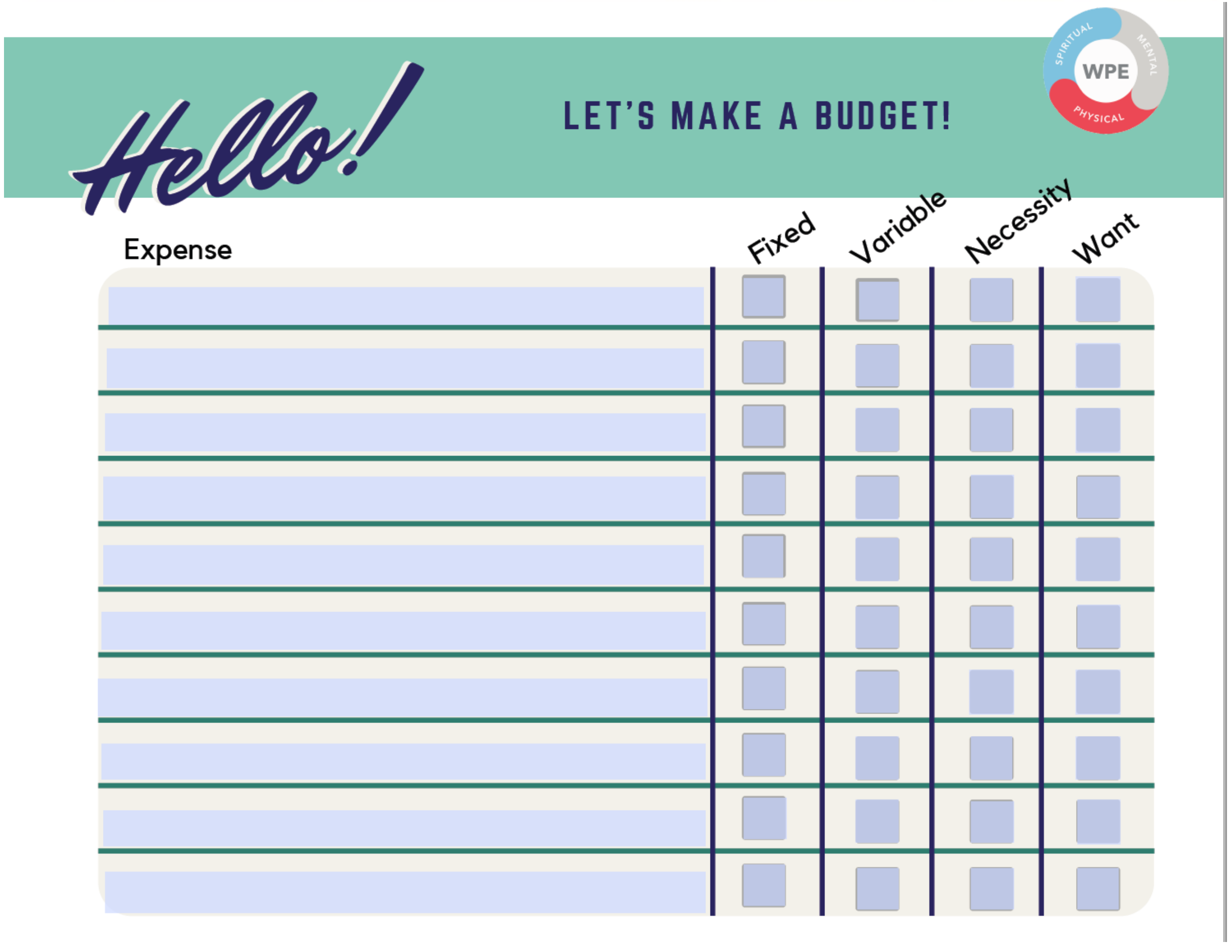


Source: California State University Channel Islands

### Post-work: thinking about credit

**Research the following questions, then discuss with either another PREP Scholar or Peer Mentor:**

- What is credit?
- What is a credit score? And how can you find yours?
- How do you build credit?
- What positively and negatively impacts credit?
- What can credit do for you or allow you to do?

### Financial Wellness

Johns Hopkins PREP Workshops

### Why care about financial wellness?

- Increases your financial security
- Enhances your freedom of choice
- Moves you from surviving to thriving each month

### Workshop objectives:

- Discussing speed budgeting exercise pre-work
- Learning more about taxes
- Heightening awareness of student loans
- Reflecting on how future goals relate to finances

### Speed budgeting exercise, discussion:

- What category of spending, if any, did you overlook?
- What are the most common ways that budgets can fail?
- Is there an aspect of your current budget you want to improve?
- How could you plan for unexpected expenses?

***—work on building an emergency fund (~3-6 months of income)***

### Tax information (federal + state):

- Yearly taxes (1 annual payment)
  - TurboTax software to file yourself
  - Tax services like H&R Block, etc.
  - W-2 form, taxes are withheld from your paycheck

- Estimated taxes (smaller payments 4x per year)
  - Applicable when taxes not withheld from paycheck—i.e., PhD program funding switch or fellowships
  - Pay online or via mail, 1040-ES form

### Universities with publicly available online tax info:
#### **be wary of state-specific info*

- Yale
- Harvard
- Northwestern
- University of Rochester

### Student loans, things to think about now:

- Who is your loan provider?
- Do you have access to your loan statements and balances?
- Which of your loans are subsidized or unsubsidized?
- Are your loans currently in deferment as a PREP Scholar?

### Student loans, things to think about for the future:

- Will your loans be in deferment as a PhD, MD or MD-PhD student?
- MDs or MD-PhDs: during medical school will you take out further federal loans, enter a loan deferment program, or apply for scholarships?
- PhDs: during graduate school will you apply for fellowships or rely on PI funding; after graduation will you enter teaching or research loan deferment programs?
- Do you have access to the FSA website?

https://studentaid.gov/

Federal Student Aid
An OFFICE of the U.S. DEPARTMENT of EDUCATION

### Future goals and finances :

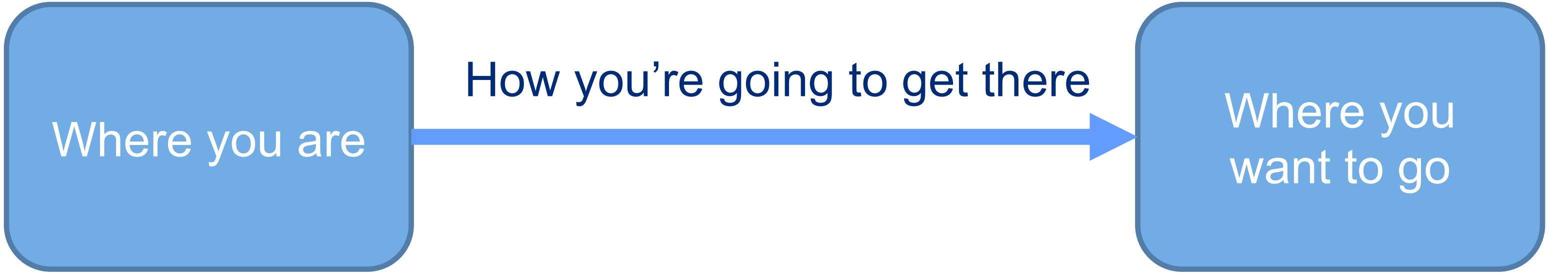


- Do your current finances align with your goals/values?
- What can you do in your current position to enhance your financial wellbeing?
- How might your finances change as you progress to your next career step?

### Application fees can be waived:

- For MSTP & PhD: apply through individual schools
  - Many schools have info and applications accessible through their websites

- For MD: AAMC Fee Assistance Program (FAP)
  - https://students-residents.aamc.org/fee-assistance-program/fee-assistance-program-fap

### Case study #1:

- Katrina (a PREP scholar) wants to help her parents pay unexpected medical bills by getting a second job. She's apprehensive about telling her current PI.
  - What advice would you give to her?
  - How could she adjust her budget to help her family while remaining financially stable herself?

### Case study #2:

- A new cohort of PREP scholars arrive in Baltimore. Robert has his heart set on living in an expensive penthouse in DC and commuting to lab each day.
  - What advice would you give him on how to make his decision?
  - What questions do you think he should be asking when finding housing?

#### Case study #3:

- PREP scholar Emmanuel is interviewing at PhD programs. His budget is tight since his apartment’s rent increased last year. His first-choice graduate schools are across the country.
  - What financial questions, in any, should he ask during his interviews?

### Links to resources:

- **Talk to PREP resources: program director, peer mentors, each other!**

- **Get a Financial Life Book (~$12 on Amazon)**
https://www.amazon.com/Get-Financial-Life-Personal-Twenties/dp/1476782385

- **Monthly budgeting calculator**
https://www.huntington.com/Personal/Calculators-Educators/budgeting-calculators/how-much-am-i-spending

- **TurboTax**
https://turbotax.intuit.com/lp/ppc/4403?srqs=null&cid=ppc_gg_b_stan_all_na_Turbo-Tax_ty21-bu2-sb5_586481380782_58623458573_kwd-11150971&srid=Cj0KCQjwmdGYBhDRARIsABmSEePELW1KjWsH0mqp7YVkYhzfyHWs0o6zumAIR0Iidx2IZeknfyDL4PcaAh0kEALw_wcB&targetid=kwd-11150971&skw=turbo%20tax&adid=586481380782&ven=gg&gclid=Cj0KCQjwmdGYBhDRARIsABmSEePELW1KjWsH0mqp7YVkYhzfyHWs0o6zumAIR0Iidx2IZeknfyDL4PcaAh0kEALw_wcB&gclsrc=aw.ds

- **H&R Block**
https://www.hrblock.com/tax-software/basic-tax-software/

- **Free way to file taxes with IRS**
https://www.irs.gov/filing/free-file-do-your-federal-taxes-for-free

### More links to resources:

- **Living wage calculator**

https://livingwage.mit.edu/

- **How to defer student loans while in grad school**

https://studentloanhero.com/featured/defer-student-loans-grad-school/

- **Medical school loan info**

https://students-residents.aamc.org/premed-navigator/5-things-i-wish-i-knew-premed-about-how-pay-medical-school

- **Medical school scholarship info**

https://www.aafp.org/students-residents/medical-students/begin-your-medical-education/debt-management/funding-options/scholarships.html

- **Free website for student discount codes at select retailers**

https://www.myunidays.com/US/en-US

- **Free website browser extension for promos and cash back at select retailers**

https://www.joinhoney.com/paypal

### Even more links to resources:

- **Teaching loan forgiveness info**

https://studentaid.gov/manage-loans/forgiveness-cancellation/teacher

- **National Institutes of Health research driven loan forgiveness program info**

https://www.lrp.nih.gov/

- **Free way to view your credit score**

https://www.creditkarma.com/

- **Various "calculators" for your money**

https://bethkobliner.com/calculators/make-home-budget/

- **General info on financial wellness**

https://www.annuity.org/personal-finance/financial-wellness/#:~:text=Financial%20wellness%20is%20a%20state%20of%20being%20in%20which%20you,U.S.%20Consumer%20Financial%20Protection%20Bureau.
