## Supplementary material for "Deepening biomedical research training: Community-Building Wellness Workshops for Post-Baccalaureate Research Education Program (PREP) Trainees": S3 Slidedeck for Identifying Stressors Workshop

### Overview

- **Goal:** The goal of this workshop is to provide tools and resources for scholars to effectively identity stressors in their life and determine best approaches for stress reduction.
- **Summary:** The workshop focused on exploring the different types of stress, the physical and emotional manifestations of stress, and discussing the nuance between healthy and unhealthy stress. Tools and resources to reduce stress were provided in the form of discussion and a resources list.

### PreWork

-Ask scholars to characterize their stress levels using music, memes, however they express themselves at different levels of stress using the template on the next slide

-Share these slides at the beginning of the workshops

Are you ready to tackle YOUR stress?

#### LET'S PLAY:

### RATE MY STRESS!

#### YOUR NAME

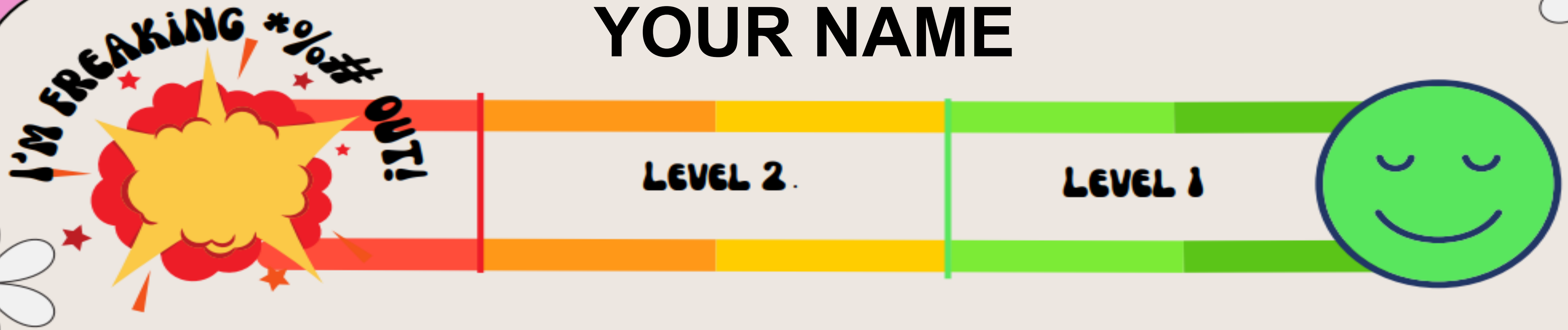


**Icebreaker**

**Activity:** Characterize your stress levels using music, memes, how ever you express yourself. What song, meme, or photo best represents your stress at Level 1, Level 2, and Level 3?

### Post-work

1. List two healthy short-term *and* long-term stress coping strategies that you'll use for "medium" and "high"-level stress.

2. Reflect back on the Values & Identity Workshop. Do your short-term and long-term stress coping strategies align with your previously stated values? Which ones?

3. List three stress management resources from our provided sources at the end of the presentation (or other resources you have identified) that you have researched/looked into.

Johns Hopkins
SCHOOL of MEDICINE

### Recognizing Stressors and Stress Management Workshop

Johns Hopkins PREP Workshops

Are you ready to tackle YOUR stress?

**LET'S PLAY:**

### RATE MY STRESS!

#### YOUR NAME

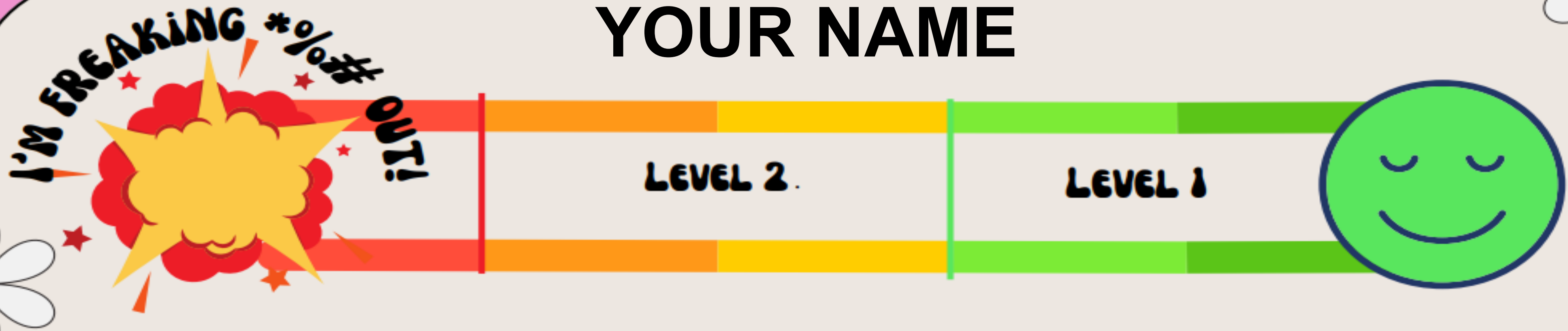


**Icebreaker**

**Activity:** Characterize your stress levels using music, memes, how ever you express yourself. What song, meme, or photo best represents your stress at Level 1, Level 2, and Level 3?

### What is stress?

### What does stress look like?

#### Physical

- Difficulty breathing
- Sleep issues
- Panic attacks
- Fatigue
- Muscle aches and headaches
- Chest pains and high blood pressure
- Indigestion
- Heartburn
- Constipation or diarrhea
- Sudden weight gain or loss
- Changes to your menstrual cycle
- Sweating

#### Emotional

- Find it hard to make decisions
- Unable to remember things
- Slow thoughts
- Constantly worrying/having feelings of dread
- Irritable
- Eating too much or too little
- Substance abuse
- Restless, like you can't sit still
- Crying or feeling tearful
- Withdrawing from people around you
- Unable to concentrate

### Types of Stress: Acute Stress

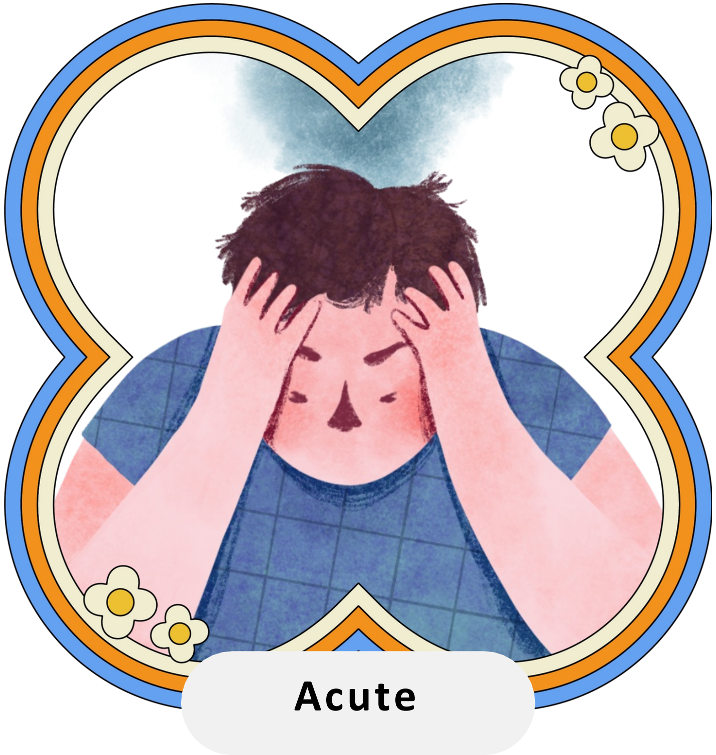


- The most common stress
- Typically brief
- Predominated by negative thoughts of situations or events in the present/future
- Causes transient physical & emotional distress

Figure made using Canva

### Types of Stress: Episodic Acute Stress

- Frequently experience acute stress
- Always feel in a rush or pressured
- Live a life of chaos & crisis
- Intense emotional, physical AND cognitive distress
- Commonly affects the following personality types: Type A & worriers

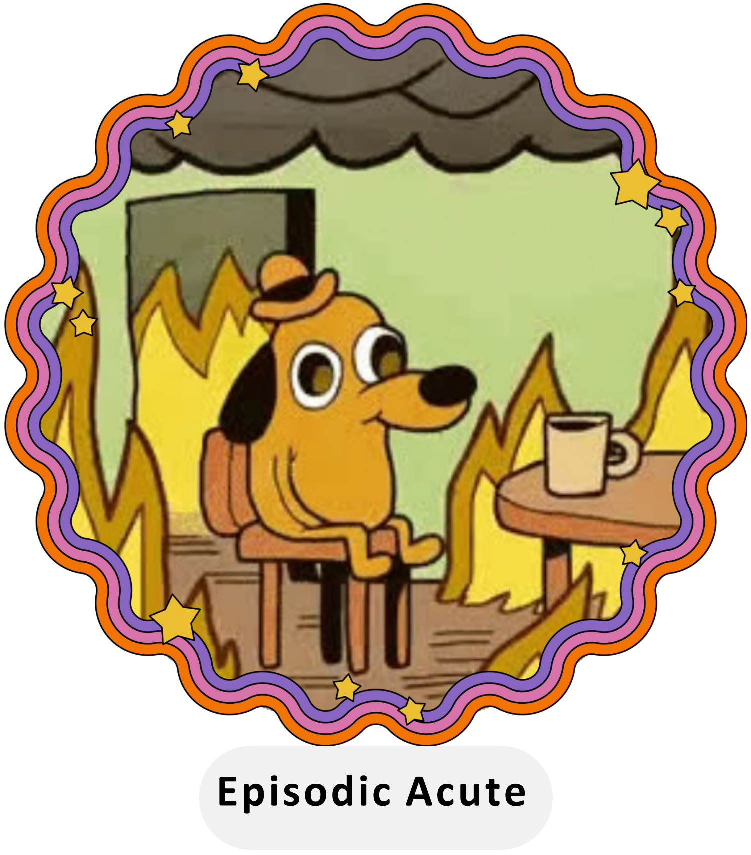


Figure made using Canva

### Types of Stress: Chronic Stress

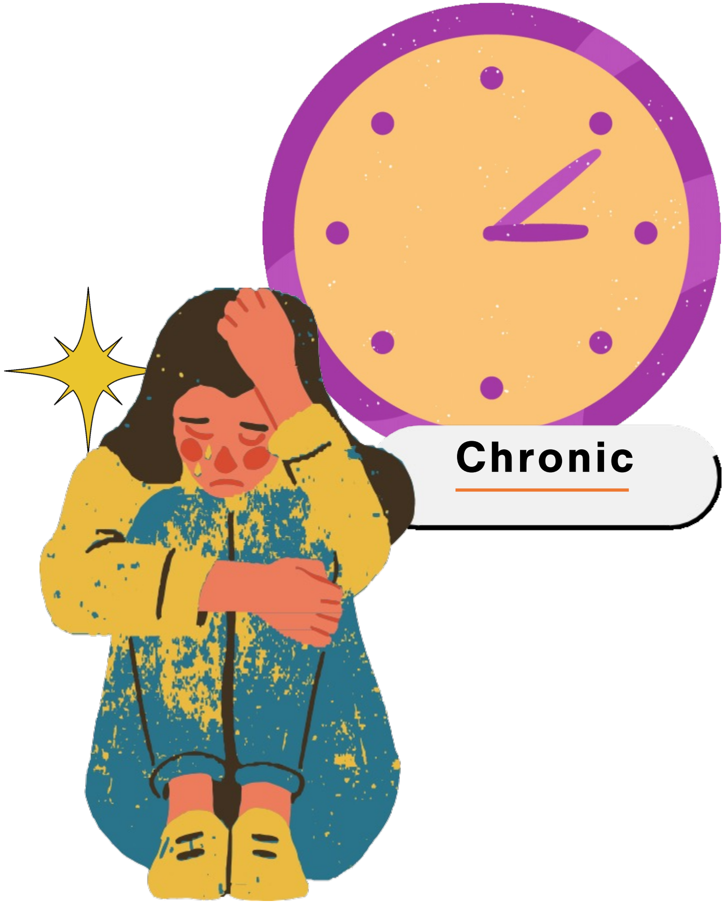


- Constant, grinding stress
- Persistent feelings of hopelessness
- Intense, crippling emotional, physical and cognitive distress
- Can be caused by childhood trauma, poverty, abuse, and long-term unhappiness

Figure made using Canva

### Name that stress case exercise: Scenario 1

Shaun has an F31 & NSF application that are both due in only a couple weeks. The thought of sitting down and finishing up both applications overwhelms him to the point that he begins sweating, hyperventilating, and feeling dizzy. He constantly feels exhausted and dreads the thought of writing/figure making anytime he thinks about the upcoming deadline.

**Acute Stress?** **Episodic Acute?** **Chronic?**

### Name that stress case exercise: Scenario 2

Shannon's relationship with her PI and lab mates has become increasingly toxic over the years. She is no longer comfortable asking for guidance or help from anyone in lab and feels isolated. She is currently a $4^{th}$ PhD student and still has several years left of her PhD before she can graduate. She constantly feels exhausted, depressed, and doesn't see any hope for a solution to her situation. She can't remember the last time she was happy.

**Acute Stress?** **Episodic Acute?** **Chronic?**

### Name that stress case exercise: Scenario 3

Collin is a 5th year PhD student who is involved in multiple student organizations, is a mentor to multiple students every academic year, and takes part-time care of a family member that lives nearby. Collin lives in a near constant state of feeling overwhelmed, anxious, and in a rush. His friends have recommended he seek professional help to build better stress management habits but he insists he can handle everything on his own and prefers to carry out tasks himself rather than delegate to others.

**Acute Stress?** **Episodic Acute?** **Chronic?**

### Healthy and Unhealthy Stressors

#### Healthy Stress

- Motivates you
- Inspires you
- Focuses your energy
- Enhances performance

#### Unhealthy Stress

- Anxiety
- Confusion
- Poor concentration
- Decreased performance

### Stressor Maintenance: Discussion

Discuss your real life examples of healthy and unhealthy stress. Identify your appropriate level of healthy stress for a lab workday. How does that change during the week? Based on your goals? What is an acceptable level of stress for you?

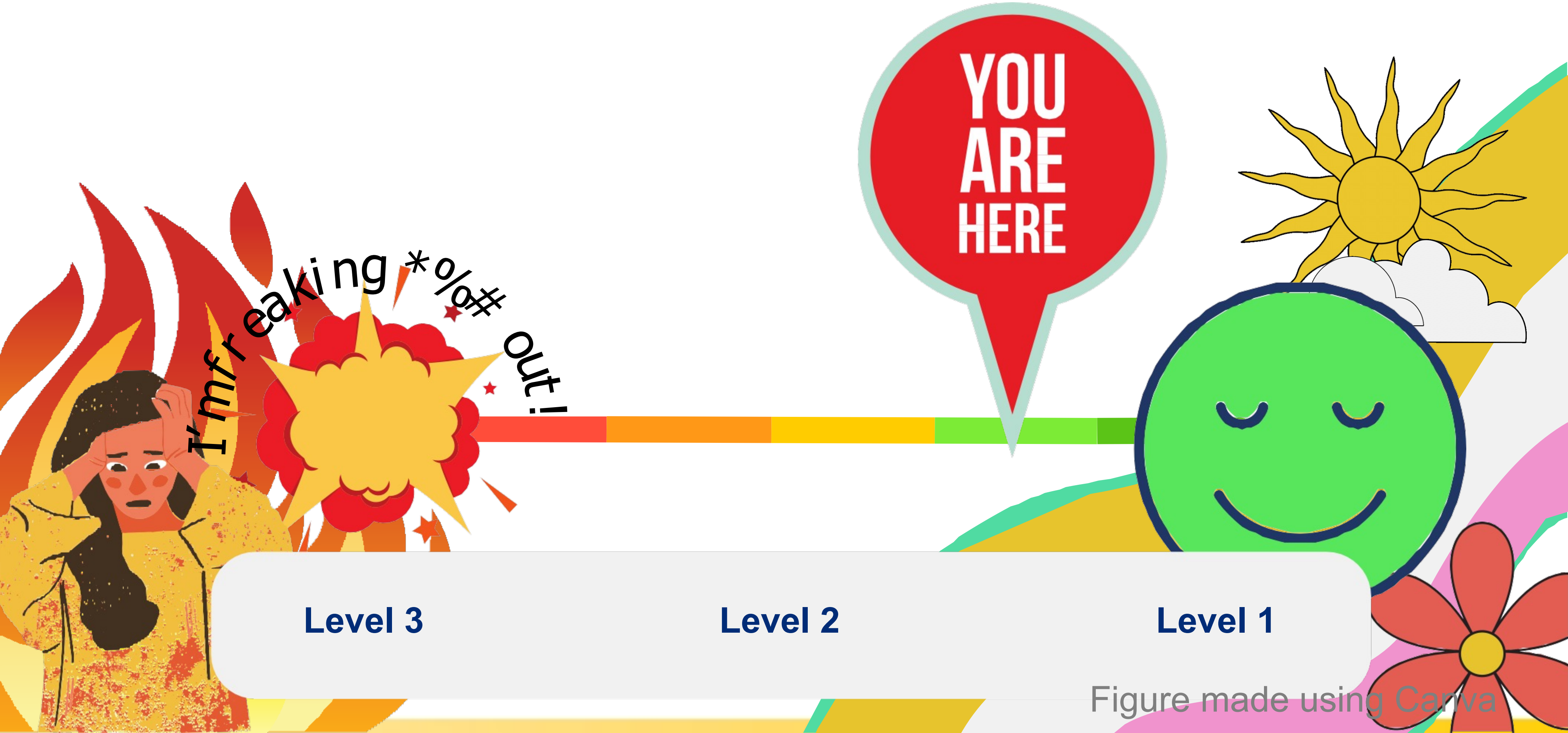


Figure made using Canva

### Identifying Stressors: Discussion and Activity

**Discussion:** What are the stressors that take you up a level? What are your behavioral and physical indicators of stress for each level? Feel free to use the chart below to guide you through the exercise.

| | Level 3 Stress | Level 2 Stress | Level 1 Stress |
|---|---|---|---|
| **My Triggers** | | | |
| **My Stims** | | | |

### Destressing: how do you destress?

- **Activity & Discussion:** What are your short-term and long-term tactics for reducing your stress?

**Short-term:**

**Example:** My slide and coverslip broke. I really needed that sample.

**Response:** Deep breathing so I don't flip this table!

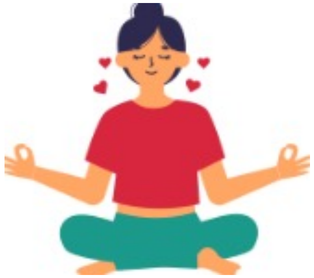

**Long-term:**

**Example:** I have been going really hard this week. I'm becoming exhausted.

**Response:** Saturdays are my day off. Every Saturday, unless dire. I will not discuss science on Saturdays.

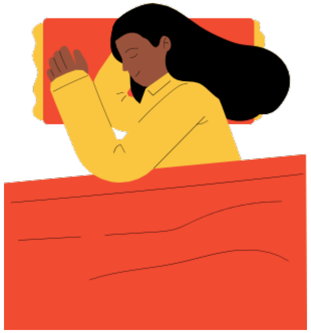

Figure made using Canva

### Things to consider: Taking it back to our Values & Goals workshop

- How have your values influenced your goals?
- Do your core values consider your mental health? Do those values extend to how you plan to achieve your goals?
- What questions do you have to ask yourself to determine whether they fit?
- For some goals, stress is unavoidable. Who will be your support network be?
- How do you think your goals align with your abilities to manage your stress?
- Do your values fit with the goals you have?
- Are your goals set in stone or are they fluid? What aspects are open to change, if any?

### Institutional Resources

- Provide a list of resources available to the students at your institution such as offices of student life, health systems, and any mental health resources

### Other Resources

#### Mental Health APPS

**Best Overall:** Moodfit

**Best for Therapy:** BetterHelp

**Best for Learning Coping Skills:** MoodMission

**Best for Stress Relief:** Sanvello

**Best for Meditation:** Calm

**Best Fun App:** Happify

**Best for Depression:** Depression CBT Self-Help Guide

**Best for BIPOC:** Shine

**Best for Bipolar Disorder:** eMoods **Best for Symptom Tracking:** Bearable

**Best for ADHD:** Todoist

**Best for PTSD:** PTSD Coach

#### Mental Health Resources

**Free online resources courtesy of:**

*Oprah Winfrey*

https://www.oprah.com/omagazine/free-online-resources-for-mental-illness

*Megan Thee Stallion https://www.badbitcheshavebaddaystoo.com/* *Nature*

https://www.nature.com/collections/gnlwffjgtr

### Sources


- *Signs and symptoms of stress*. Mind. (n.d.). https://www.mind.org.uk/information-support/types-of-mental-health-problems/stress/signs-and-symptoms-of-stress/
- Shawna Freshwater, PhD. (2020, January 18). *3 types of stress and health hazards*. Shawna Freshwater, PhD. https://spacioustherapy.com/3-types-stress-health-hazards/
- *Stress management: How to tell the difference between good and bad stress*. Summa Health. (n.d.). https://www.summahealth.org/flourish/entries/2021/01/stress-management-how-to-tell-the-difference-between-good-and-bad-stress
- Molnar, H. (2022, November 21). *Student support: The Johns Hopkins University School of Medicine*. Johns Hopkins Medicine. https://www.hopkinsmedicine.org/som/life-at-hopkins/student-support.html
