## Supplementary material for "Deepening biomedical research training: Community-Building Wellness Workshops for Post-Baccalaureate Research Education Program (PREP) Trainees": S4 Slidedeck for Mental Health Strategies Workshop

### Overview

- In this presentation we wanted to demystify taboos about mental health by defining and highlighting the importance of mental health.
- Scholars learn about signs and symptoms of anxiety, depression, and stress and were given tools to alleviate these signs and symptoms.
- Resources offered by your local school should be included at the end.

### PreWork

JOHNS HOPKINS SCHOOL of MEDICINE

- Read: N. Forester. (2021) Mental Health of Graduate Students is sorely overlooked. Nature 595: 135-137.
- Answer the following question:  What are some way a student could manage their mental health?

### Postwork

- Next time you feel your stress levels rising, use the  Self Care Checkup and Daily Mood Tracker Worksheets (found in Supplemental Information) to track your mood and gauge the efficacy of your mental health practices.

### Mental Health Strategies

Johns Hopkins PREP Workshops

### Mental Health

- Mental health is a crucial aspect of overall well-being and is essential for leading a fulfilling life.
- Good mental health helps us to manage stress, form and maintain healthy relationships, and make meaningful contributions to our communities.

### A Response To Life’s Challenges

JOHNS HOPKINS
SCHOOL *of* MEDICINE

- Mental health refers to a person's ability to cope with life's challenges and difficulties, to manage their emotions, and to maintain healthy relationships.
- It encompasses a range of skills and abilities, including the ability to:
- Manage feelings and emotions, such as anger, sadness, and anxiety
- Form and maintain healthy relationships with others
- Adapt to change and cope with stress
- Set and achieve meaningful goals
- Contribute positively to one's community and society

### Cognitive Distortions

- Irrational thought pattern that causes the person to perceive reality inaccurately
- Common in people suffering from anxiety and depression
  - Fortune telling
  - Mindreading
  - All-or-nothing thinking
  - Jumping to conclusions
  - Overgeneralization
  - Discounting the positive, etc.

### Manifestations Of Anxiety

JOHNS HOPKINS
SCHOOL of MEDICINE

- Restlessness or feeling on edge
- Difficulty concentrating or feeling easily distracted
- Irritability or agitation
- Muscle tension or headaches
- Excessive worry or fear
- What to do when we are anxious?
  - Exercise, sleep, and eat well
  - Avoid caffeine, alcohol, and nicotine
  - Grounding techniques

### Manifestations Of Depression

JOHNS HOPKINS
SCHOOL *of* MEDICINE

- Persistent feelings of sadness or hopelessness
- Loss of interest or pleasure in activities
- Changes in appetite or sleep patterns
- Fatigue or low energy
- Thoughts of self-harm or suicide
- What to do when we are feeling down?
  - Do not self-isolate!
  - Do things that make you happy!

### Manifestations of Stress

- Increased heart rate or sweating
- Shallow breathing or hyperventilation
- Racing thoughts or difficulty focusing
- Muscle tension or aches
- Digestive problems or nausea
- How to manage stress?
  - Take a break from the news and social media
  - Make time for yourself (meditate, do yoga, painting, etc)

### Self-awareness

- Mental hygiene refers to the practice of taking care of your mental health, much like how we practice personal hygiene to take care of our physical health.
- Just as we brush our teeth, exercise, and eat well to maintain physical health, there are also practices and habits we can adopt to promote good mental health.
- Mental hygiene involves self-awareness, self-care, and seeking support when necessary.

### Self-care

- Self-care involves taking intentional actions to promote mental health and well-being.
- Self-care practices can include exercise, getting enough sleep, eating nutritious food, engaging in hobbies, and spending time with loved ones.
- Self-care also involves setting boundaries and saying no to things that don't serve our mental health.

### Seeking Support

JOHNS HOPKINS SCHOOL of MEDICINE

- Seeking support is an important part of mental hygiene. Just as we see a doctor for physical health concerns, we can seek professional help for mental health concerns.
- This can include therapy, medication, or support groups.
- Asking for help can be difficult, but it's a brave and necessary step towards better mental health.

### Recognizing when to seek support

- Recognizing when to seek support is an important part of maintaining good mental health.
- Some signs that it might be time to seek support include feeling overwhelmed, experiencing changes in mood or behavior, or having difficulty concentrating or coping with stressors.
- It's important to remember that seeking support is a sign of strength, not weakness.

### Types Of Support

- There are many types of support available for those experiencing mental health challenges.
- This can include psychotherapy, pharmacotherapy, support groups, and self-help resources.
- It's important to find the type of support that works best for you and your unique needs.

### Seeking Professional Help

- Seeking professional help, such as therapy or medication, can be a helpful part of managing mental health concerns.
- A mental health professional can provide a safe and non-judgmental space to explore thoughts, feelings, and behaviors.

### Case Study #1

- **Jenny** is a post-baccalaureate student preparing to attend medical school. She's been feeling extremely overwhelmed and anxious lately. She's been having trouble sleeping, has lost her appetite, and has been feeling fatigued all the time. She's also been avoiding her friends and family and feels like she can't keep up with the demands of her coursework.
    - What are some symptoms of poor mental hygiene that Jenny is experiencing?
    - What are some self-care strategies that Jenny can use to manage her stress and anxiety?
    - How can Jenny seek support from friends, family, or mental health professionals?
    - What are some potential barriers that may prevent Jenny from seeking help, and how can she overcome them?

### Case Study #2



- **Mike** is a graduate student in Neuroscience who has noticed that his friend **John** has been struggling with depression. **John** has been feeling very down lately and has stopped coming to class and Graduate Student Association social events. **Mike** is worried about **John's** well-being and wants to help**.**
  - What are some signs that John may be experiencing depression?
  - What are some ways that Mike can approach John and offer support without being intrusive or pushy?
  - What are some resources that Mike can recommend to John, such as counseling or mental health services?
  - How can Mike continue to support John in the long term, while also setting boundaries and taking care of his own mental health?

### Case Study #3

JOHNS HOPKINS
SCHOOL of MEDICINE

- You're a graduate student working in a lab with several other members, including a colleague who has been “working from home”. At first, they asked for little favors, such as feeding cells and/or collecting data. However, these requests have now escalated to full-day experiments, which have impacted your productivity and ability to work. You're unsure how to handle the situation.

    - How can you balance being a helpful colleague with protecting your own time and productivity?
    - What are some potential consequences of not setting boundaries in this situation?
    - How can you communicate your boundaries in a clear and respectful way?
    - If your lab mate is resistant to your boundaries, what are some potential next steps you can take?

### Discussion

JOHNS HOPKINS SCHOOL of MEDICINE

- In order to continue to work effectively and efficiently, mental health must be maintained as well as possible to stay positive and give the best result. Some of these methods and tips can help maintain your mental health.

### Step #1: Talk about your feelings

- Someone who experiences mental health instability usually always accumulates feelings of stress without expressing it. It's good to try to express and talk about your feelings so that there is no emotional outburst that can have bad consequences.

### Step #2: Eat well

- What we eat can affect how we feel both immediately and in the longer term. A diet that is good for your physical health is also good for your mental health. Therefore, you have to pay attention to what food you eat.

### Step #3: Keep in touch

JOHNS HOPKINS
SCHOOL of MEDICINE

- Relationships are key to our mental health. Working in a supportive team is hugely important for our mental health at work. We don’t always have a choice about whom we work with, and if we don’t get on with managers, work peers, or clients, it can create tension. As long as we keep good communication, it can minimize mental issues in the long run.

### Highlight local resources available at your institution
