## Supplemental Data 1 for "Deepening biomedical research training: Community-Building Wellness Workshops for Post-Baccalaureate Research Education Program (PREP) Trainees"

### Self-Care Checkup

Self-care activities help us enhance our well-being and maintain good mental health.

They can include habitual, routine activities such as eating well and getting regular exercise, which often get neglected during particularly difficult or stressful periods in our lives.

Becoming aware of how often, or how well, we practice self-care activities can help us identify areas we are neglecting and improve upon them for better mental health.

#### Instructions

This *Self-Care Checkup* can help you consider the frequency and quality of your self-care in five important life domains:

- **Emotional**
- **Physical**
- **Social**
- **Professional;** and
- **Spiritual self-care.**

Using the key provided below, rate how well, or how frequently, you believe that you engage in each activity between your therapy sessions.

It's important to remember the list is not exhaustive - some activities may not appeal to you at all, or you may feel that others are missing.

If you think of ideas that you'd like to add to the list, use the spaces provided.

| | | |
|---|---|---|
| **1** | *I rarely do this* | *I don't do this well* |
| **2** | *I sometimes do this* | *I'm average at doing this* |
| **3** | *I do this often* | *I do this very well* |
| ☐ | *I'd like to do this more often* | *I'd like to become better at this* |

| Emotional Self-Care | | |
|---|---|---|
| 1 2 3 | ☐ | Enjoying hobbies |
| 1 2 3 | ☐ | ‘Unplugging’ from technology (e.g. email, social media) |
| 1 2 3 | ☐ | Expressing emotions and feelings (e.g. talking, journaling) |
| 1 2 3 | ☐ | Appreciating own talents, accomplishments, and strengths |
| 1 2 3 | ☐ | Taking days off/rest days from responsibilities |
| 1 2 3 | ☐ | Learning about or exploring new things (e.g. hobbies, foreign languages) |
| 1 2 3 | ☐ | Practicing self-nurturing activities (e.g. long bath, gentle walk) |
| 1 2 3 | ☐ | Laughing about things |
| 1 2 3 | ☐ | Taking a holiday, escape, or mini-break |
| 1 2 3 | ☐ | General emotional self-care |
| 1 2 3 | ☐ | |

| Physical Self-Care | | |
|---|---|---|
| 1 2 3 | ☐ | Attending health upkeep appointments (e.g. dental or GP checkups) |
| 1 2 3 | ☐ | Resting when unwell |
| 1 2 3 | ☐ | Drinking enough water |
| 1 2 3 | ☐ | Getting sufficient sleep |
| 1 2 3 | ☐ | Enjoying group exercise (e.g. gym classes, hobbies) |
| 1 2 3 | ☐ | Eating regular meals |
| 1 2 3 | ☐ | Exercising out of doors |
| 1 2 3 | ☐ | Maintaining good hygiene |
| 1 2 3 | ☐ | Eating a healthy diet |
| 1 2 3 | ☐ | General physical self-care |
| 1 2 3 | ☐ | |

| Social Self-Care | |
|---|---|
| 1 2 3 ☐ | Making time for friends or family |
| 1 2 3 ☐ | Staying in contact with distant connections (e.g. Skype, Facetime) |
| 1 2 3 ☐ | Engaging in mentally stimulating discussions |
| 1 2 3 ☐ | Being intimate/romantic with partner |
| 1 2 3 ☐ | Asking for help when you require it |
| 1 2 3 ☐ | Doing fun activities with others/ enjoyable group activities |
| 1 2 3 ☐ | Spending quiet private time with partner |
| 1 2 3 ☐ | Making new friends/talking to new people |
| 1 2 3 ☐ | Overall social self-care |
| 1 2 3 ☐ | |

| Professional Self-Care | |
|---|---|
| 1 2 3 ☐ | Seeking support when it's required at work |
| 1 2 3 ☐ | Maintaining a comfortable or pleasant work environment |
| 1 2 3 ☐ | Socializing or bonding with co-workers |
| 1 2 3 ☐ | Balancing work and leisure activities |
| 1 2 3 ☐ | Accepting stimulating/interesting new tasks or projects |
| 1 2 3 ☐ | Taking lunch breaks/regular work breaks |
| 1 2 3 ☐ | Turning down unnecessary/ unreasonable tasks |
| 1 2 3 ☐ | Pursuing further professional development opportunities |
| 1 2 3 ☐ | Seeking recognition/promotion/reward where deserved |
| 1 2 3 ☐ | General professional self-care |
| 1 2 3 ☐ | |

| Spiritual Self-Care | | |
|---|---|---|
| 1 2 3 | ☐ | Enjoying outdoor/nature time |
| 1 2 3 | ☐ | Volunteering for charity/community |
| 1 2 3 | ☐ | Religious practice |
| 1 2 3 | ☐ | Practicing gratitude |
| 1 2 3 | ☐ | Meditating |
| 1 2 3 | ☐ | Allocating quiet time for reflection |
| 1 2 3 | ☐ | Applying personal strengths, talents, or values |
| 1 2 3 | ☐ | Appreciating beauty (e.g. music, art, literature) |
| 1 2 3 | ☐ | General spiritual self-care |
| 1 2 3 | ☐ | |
