## Supplemental Data 2 for "Deepening biomedical research training: Community-Building Wellness Workshops for Post-Baccalaureate Research Education Program (PREP) Trainees"

### Daily Mood Tracker

| | Happy | Sad | Angry | Excited | Anxious | Tired | Other | Notes |
|---|---|---|---|---|---|---|---|---|
| 06:00 – 08:00 | ☐ | ☐ | ☐ | ☐ | ☐ | ☐ | ☐ | |
| 08:00 – 10:00 | ☐ | ☐ | ☐ | ☐ | ☐ | ☐ | ☐ | |
| 10:00 – 12:00 | ☐ | ☐ | ☐ | ☐ | ☐ | ☐ | ☐ | |
| 12:00 – 14:00 | ☐ | ☐ | ☐ | ☐ | ☐ | ☐ | ☐ | |
| 14:00 – 16:00 | ☐ | ☐ | ☐ | ☐ | ☐ | ☐ | ☐ | |
| 16:00 – 18:00 | ☐ | ☐ | ☐ | ☐ | ☐ | ☐ | ☐ | |
| 18:00 – 20:00 | ☐ | ☐ | ☐ | ☐ | ☐ | ☐ | ☐ | |
| 20:00 – 22:00 | ☐ | ☐ | ☐ | ☐ | ☐ | ☐ | ☐ | |
| 22:00 – 00:00 | ☐ | ☐ | ☐ | ☐ | ☐ | ☐ | ☐ | |
