## Supplementary material for "Deepening biomedical research training: Community-Building Wellness Workshops for Post-Baccalaureate Research Education Program (PREP) Trainees": S7 Slidedeck for Self Advocacy Workshop

### Overview

- The goal of this workshop is to promote self-advocacy to PREP scholars by providing examples of different communication styles and raising awareness for boundary setting
- Although assertive communication will be promoted, we will also provide examples when passive or aggressive communication could be used

### Pre-work: What is your communication style?

**Briefly reflect on the following questions and be prepared to share your response if you feel comfortable:**

- How would you describe your general communication style?
- How similar/different is it to the communication style that is modeled by your family/friends/peers/and mentors?

### Post-work: Reflect on your communication style

- Think about an important relationship at work or at home. Consider a time when you struggled to be assertive and communicate your needs effectively. Focus on what happened, why it happened, how it made you feel, and anything you wish had gone differently.

- Now focus on a time you were assertive and communicated your needs effectively. Focus on what happened, why it happened, how it made you feel, and anything you wish had gone differently.

### Self-Advocacy

Johns Hopkins PREP Workshops

JOHNS HOPKINS
SCHOOL *of* MEDICINE

### Workshop objectives:

- To identify different styles of communication including passive, passive-aggressive, aggressive and assertive

- To define self-advocacy and context-dependent boundary setting

- To obtain tools to help with assertive communication

### Introduction Question:

- What is self-advocacy?
- What is assertiveness?
- How do these two differ?

### Two Important Definitions

- Self-advocacy is…
  - The action of representing ourself and our views/interests with the goal of meeting our needs
  - One or two types of advocacy – for ourself and for others; for many of us it is easier to advocate for others
- Assertiveness is the ability to…
  - Express one's feeling and assert one's rights and needs while respecting the feelings, rights and needs of others
  - Use communication that is direct, open and honest to address situations that concern you
  - Set appropriate boundaries that feel right given the context and situation

### Another Way to Look at These Definitions

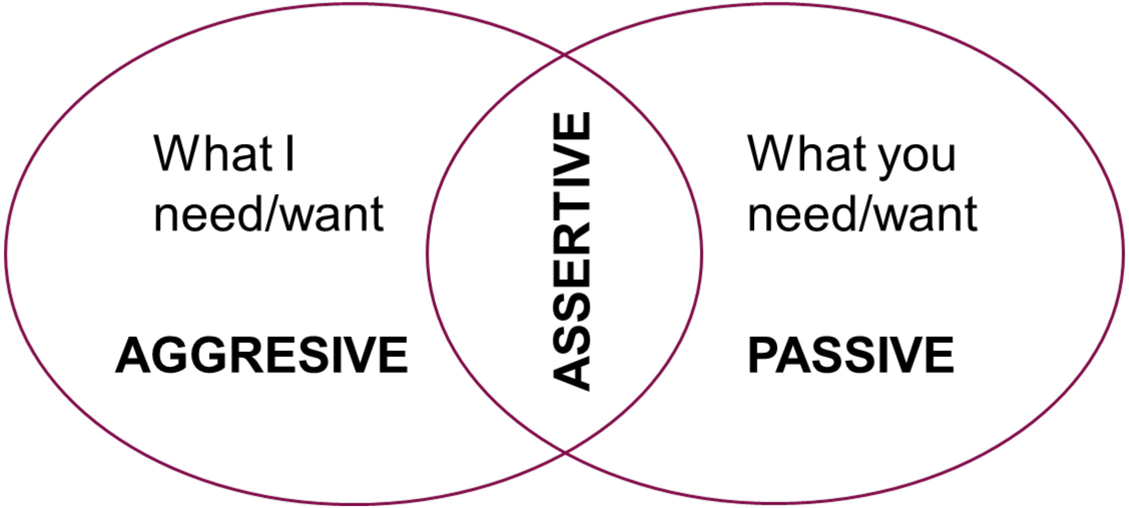


**RESPECT and BOUNDARIES are key elements of assertive communication**

Figure courtesy, Dr. Sharon L. Milgram, Director NIH OITE

### Identifying Different Communication Styles

- Passive Communication – Unexpressive; "what happens, happens."
- Aggressive Communication – Hostile expression of one's own needs
- Passive-aggressive Communication – Behind the scenes anger

### The Passive Communication Style

- Does not directly express feelings, needs and wants and prefers to avoid conflict and confrontation
- Can have build-up feelings of resentment and anger, which can lead to an anger outburst
- Outcomes:
  - Can lead to feelings of anxiety/depression and feeling out of control
  - Prone to feeling confused due to ignoring their own feelings
- **EXAMPLE: Your PI asks you to take on a summer student during a busy season full of experiments. Your reply: "Sure, if you think that would be a good idea."**

### The Aggressive Communication Style

- Directly expresses feelings, needs and wants without considering others
- Prone to criticizing, blaming and can be intimidating
- May not listen as well and is defensive or hostile when confronted by others
- Outcomes:
  - Alienates others and instills resentment in them
- **EXAMPLE: Your summer student says they cannot come to lab on Tuesday mornings due to their summer programming. You reply: "If you want to have a successful summer, I expect you to make up those hours to help finish those experiments."**

### The Passive-Aggressive Communication Style

- Does not directly express feelings, needs and wants but attempts to do so in a subtle manner that may convey frustration – resorts to sarcasm
- Denies conflict by exhibiting avoidant behavior and may feel powerless and resentful
- May show negative feelings through actions or attitude
- Outcomes:
  - Alienates others
  - May feel stuck in your own situation
- **EXAMPLE: Your summer student accidently misloads a Western Blot that you planned for them to do. Now you do not know the order of the samples in the lanes. Your reply: "This sucks. Maybe if you were here more often you would've known the right loading order of the samples."**

### The Assertive Communication Style

- Directly expresses feelings, needs and wants - use lots of “I” statements
- Have relaxed and confident body language
- Good-listeners and do not interrupt
- Outcome:
  - Oriented towards growth because they address issues as they come up
  - Maintain connections to others
- **EXAMPLE: Your PI asks you to take on a summer student during a busy season full of experiments. Your reply: "I have many important experiments planned this season. I do not think I will have time to work full time with a summer student. However, I would like to help with whatever spare time I have. Maybe I can share the responsibility with another grad student?"**

### Situations That Call For Different Communication Styles

- Passive Communication
  - A viable option if conflict can lead to an escalation of violence
- Aggressive Communication
  - If someone is in imminent harm and you must act and communicate fast to remove them from that situation

### Additional Notes on Communication Styles

- We may not always resort to a single communication style in all our relationships but display a mix of many depending on the circumstances
- There are even different boundaries of what passive and aggressive communication style look like that can intersect with identity.
  - For example, to individuals of different cultures, what appears as assertive communication can come across as aggressive or what is assertive as passive
  - Women or individuals from minority groups displaying assertive communication may be seen as "aggressive"

### Scenario #1:

"You are working on a project with the guidance of an assistant professor that is directly under your PI. You have finished acquiring and analyzing the data for this project, and you give a brief presentation to update your PI and the assistant professor on the progress that has been made. Your PI replies: 'Great job! Someone should start writing the paper.' Unknown to you, the assistant professor has begun to write the manuscript. You feel that you have been denied a key learning opportunity of being an active participant in writing the manuscript."

- **In this situation...**
  - **What advice would you give to the person in this situation?**
  - **What would you say if you were in this situation?**

### Scenario #2:

"Your PI is applying for research funding in a field that is new but promising for your lab. To generate preliminary data quickly, your PI has designed a few pilot experiments that will take at most 4 weeks to accomplish. Your PI has asked you to work with a colleague to get these pilot experiments done. You recognize that you don't work very well with this colleague because they work at a different pace from you and take credit for your contributions without putting much effort in on their part. In this case, you think that it might be easier if only one person performs the pilot experiments"

- **In this situation...**
    - **What advice would you give to the person in this situation?**
    - **What would you say if you were in this situation?**

### Scenario #3:

"You and a colleague have a report due at the end of the week. You also have another project and are very busy. You meet with your colleague to go over the report and the two of you realize you need to redo some of the sections and add others. Your colleague tells you they cannot do much because they have family visiting this whole week."

- **In this situation...**
  - **What advice would you give to the person in this situation?**
  - **What would you say if you were in this situation?**

### Tips for Striving Towards Assertive Communication

- Consider your values and the risks and rewards of your decisions and its impact on your relationship.
- Use "I" statements to avoid sounding accusatory.
- Practice saying "no" with a brief explanation when appropriate.
- Rehearse with family and friends situations that you have encountered or may encounter. Ask for feedback.
- Practice assertive body language even if you don't feel it. Try to maintain a neutral or positive tone, maintain eye contact and sit or stand upright.
- If you need time or a break from conflict, do so until you can try to get your emotions in check. Remind yourself that it's okay if you make "mistakes" while learning assertiveness.
- Remind yourself that this is a process. You can strive towards being assertive without feeling fully confident in your ability towards it. Try to come up with phrases that activate you towards assertive communication.
