## Supplementary material for "Deepening biomedical research training: Community-Building Wellness Workshops for Post-Baccalaureate Research Education Program (PREP) Trainees": S8 Slidedeck for Racial Bias and Imposter Syndrome

### Overview

- The goal of this workshop is for scholars to understand the relationship between racism and imposter syndrome, to reflect on self-affirmations, and use peer coaching as a way to develop their own solutions.
- It is critical at the start of this workshop to establish this event as a safe space and acknowledge the potential triggers in this topic.

### Prework – Part 1

- Ask scholars to reflect on what you do/use for self-affirmation.
- Read this article regarding how discrimination feeds imposterism.
  - "The legacies of systemic and internalized oppression: Experiences of microaggressions, imposter phenomenon, and stereotype threat on historically marginalized groups" (2021) Nadal et al. (2006) New Ideas in Psychology.

### Prework – Part 2

- Further optional reading
  - **Johnson, V. E., Nadal, K. L., Sissoko, D. R. G. & King, R. “It’s Not in Your Head”: Gaslighting, ‘Splaining, Victim Blaming, and Other Harmful Reactions to Microaggressions. *Perspect. Psychol. Sci.* 16, 1024–1036 (2021).**

### Post-work

- **•For the follow up activity there will be two parts using the Peer Coaching Document adapted from resources provided by the BRAINS Program (**Yen, J. W., Horner-Devine, M. C., Margherio, C. & Mizumori, S. J. Y. The BRAINS Program: Transforming Career Development to Advance Diversity and Equity in Neuroscience. *Neuron* **94**, 426–430 (2017). & https://blogs.biomedcentral.com/bmcblog/2017/05/18/peer-mentoring-circles-a-strategy-for-thriving-in-science/ )
  - **#1 With your same partner from the workshop begin by swapping roles. Near peer mentors that previously asked questions will now present a dilemma statement (This can include things that cause imposterism or other dilemmas), be questioned by your partner and produce an action plan at the end of questions.**
  - **#2 Afterwards, swap back roles and have a follow-up session**

### Imposter Syndrome

Johns Hopkins PREP Workshops

### What one word immediately comes to mind when you think of Imposter Syndrome?

Gather responses from the group

**“I often feel like I am not good enough for my position, and that one day I will be discovered to be a fraud.”**

### What is Imposter Syndrome?

- ❑ "Described as individual's intellectual self-doubt and fear of failure, which is characterized by concern that others have overestimated their talents or abilities and therefore feel defrauded" (*1*).

- ❑ "This doubt can negatively impact educational persistence, achievement, and mental wellness (*2*).

1. P. R. Clance, S. A. Imes, The imposter phenomenon in high achieving women: Dynamics and therapeutic intervention. *Psychotherapy Theory Res Pract*. 15, 241–247 (1978).
2. S. Craddock, M. Birnbaum, K. Rodriguez, C. Cobb, S. Zeeh, Doctoral Students and the Impostor Phenomenon: Am I Smart Enough to Be Here? *J Student Aff Res Pract*. 48, 429–442 (2011).

### Common Symptoms Attributed to Imposter Syndrome

- ❑ Sees own accomplishments due to luck
- ❑ Feel like you are never enough
- ❑ Attributes failures to self-worth
- ❑ Practices negative self-talk
- ❑ Fear of being discovered as a fraud

### Impact of Imposter Syndrome

- Inability to give self grace
- Obsesses over flaws
- Self-sabotages by: overworking, refusing help, procrastinating

#### Causes of Imposter Syndrome

- ❑ Internal self-doubt

- ❑ Discrimination!!

### Group reflection on your own experiences with imposter syndrome.

### Discrimination and Imposterism

**“While imposter syndrome is real, internalized imposter syndrome is often misconstrued to be the root of self-doubts of those underrepresented in science without investigating structural issues in academia. Often nonbelonging rooted in discrimination (racism, genderism, heterosexism, xenophobia, ableism, ageism, classism, etc.) (3).”**

3. King, R. Microaggressions, Imposter Phenomenon, and People of Color: A Quantitative Analysis.

### Discrimination and Imposterism (cont.)

**“The present conceptualization of imposter phenomenon is problematic because it tends to focus only on psychological process associated with imposterism while generally failing to consider how the processes are shaped by interactions and structures…… And since these structural conditions create a host of negative outcomes, usage of the term ‘imposter syndrome’ ignores and grossly minimizes institutional factors, policies, and practices that cause Black (and other marginalized) students to logically respond with distress and frustration (*4*).”**

4. E. O. McGee, P. K. Botchway, D. E. Naphan-Kingery, A. J. Brockman, S. Houston, D. T. White, Racism camouflaged as impostorism and the impact on Black STEM doctoral students. Race Ethn Educ. 25, 487–507 (2022).

### Contributions of discrimination to imposterism.

**Deidentified first-hand accounts.**
**Discuss how each of these accounts may impact sense of belonging and imposterism.**

**“One day I was having coffee and other delicious treats with a group of black women at the coffee shop on campus, which we do actually once every month. When the waiter brought us the check, he said, ‘Do not leave without paying.’”**

When my school was asked by the dean to come up with a plan to help the Baltimore community after the events that surrounded Freddie Gray's death, I was specifically called upon by a departmental chair to help come up with ideas on what our department could specifically do. I was later called into a faculty meeting to express these ideas (which I never expected to have to do), which showed me that instead of focusing on being a student and earning my PhD, I have become a spokesperson for my race.

**“The reason you got into that lab is because they needed diversity points.”**

### Group reflection on ideas to navigate non-belonging whether due to discrimination or imposter syndrome

### Battling Discrimination and Imposterism

- ❑ Don’t compare your insides to someone else’s outsides.
  - ❑ Write down and celebrate your small wins!
- ❑ Journeys are different. Your timeline is your own.
- ❑ Seek out mentors, sponsors, and advocates that will openly call out discrimination and are willing to have conversations to support your needs.
- ❑ Build community!- share structured examples that exist within and outside of Hopkins.
- ❑ Start/participate a peer coaching circle!- a good way to investigate your thoughts and think of professional solutions with peers.

### Peer Coaching Group Activity

**Dilemma statement: I feel [emotion] about [situation] and I want to [goal].**

**Sample coaching questions:**
What would success look like?
What would you like to see or experience moving forward?
What have you tried so far?
What support do you want to move forward?
How do you like to make decisions?
What do you need to know to make a decision?
What has worked for you in the past?

https://blogs.biomedcentral.com/bmcblog/2017/05/18/peer-mentoring-circles-a-strategy-for-thriving-in-science/

### Peer Coaching Group Activity (cont.)

**Peer Coaching Circle (PCC) NORMS AND COMMITMENTS:** the PCC experience is grounded in a commitment to community norms and framework. We share some of these here and will touch on them again when we meet

- *Confidentiality*. Everything said and heard in each PCC conversation will remain confidential.

- *Commitment*. Each PCC participant will commit to meeting regularly and actively participate in the group process for at least 6 months.

- *Listening*. Each PCC participant will listen without judgment and with an open mind in order to reflect back what they are hearing and create opportunities for individuals to find their own best solutions.

https://blogs.biomedcentral.com/bmcblog/2017/05/18/peer-mentoring-circles-a-strategy-for-thriving-in-science/

### Practice Peer Coaching in Pairs

- ❑ 2 minutes to make dilemma statement
- ❑ 8 minutes of questions
- ❑ 2 minutes to form contract

https://blogs.biomedcentral.com/bmcblog/2017/05/18/peer-mentoring-circles-a-strategy-for-thriving-in-science/
