## Supplementary material for "Deepening biomedical research training: Community-Building Wellness Workshops for Post-Baccalaureate Research Education Program (PREP) Trainees": S9 Slidedeck for Difficult Conversations

### Overview

- The goal for the Difficult Conversations workshop was to define situations specific to biomedical sciences research that may warrant a difficult conversation and share various tools and frameworks for having this type of conversation. Along with other professions, different conflicts arise while working in biomedical research that span minor annoyances to major disputes. Understanding how to evaluate these varying conflicts and planning for how to resolve these conflicts are key tenets to maintaining healthy and effective working relationships with others.

### Pre-work

- Scholars should think about the following questions in preparation for the Difficult Conversations workshop:

- Think of a time you had an interaction in the lab/workplace that made you feel hurt, uncomfortable, or disrupted your overall mental health. What was the situation? How did it make you feel? Was a difficult conversation necessary to resolve the situation?

### Post-work

- Please review the handouts entitled "Courageous Conversations Discussion Planner" and "Courageous Conversations Reflection Worksheet". The first handout details how to have a difficult conversation similar to the steps covered in this workshop and the second is a useful worksheet to help you identify the conflict and how to address it. We encourage you all to use these and practice filling out the worksheet to help navigate how to have difficult conversations.

Worksheets from Annalisa Peterson, Ombuds for Doctoral and Postdoctoral Students, Fellows and Programs, Johns Hopkins University

### Difficult Conversations

Johns Hopkins PREP Workshops

### What is a difficult conversation?

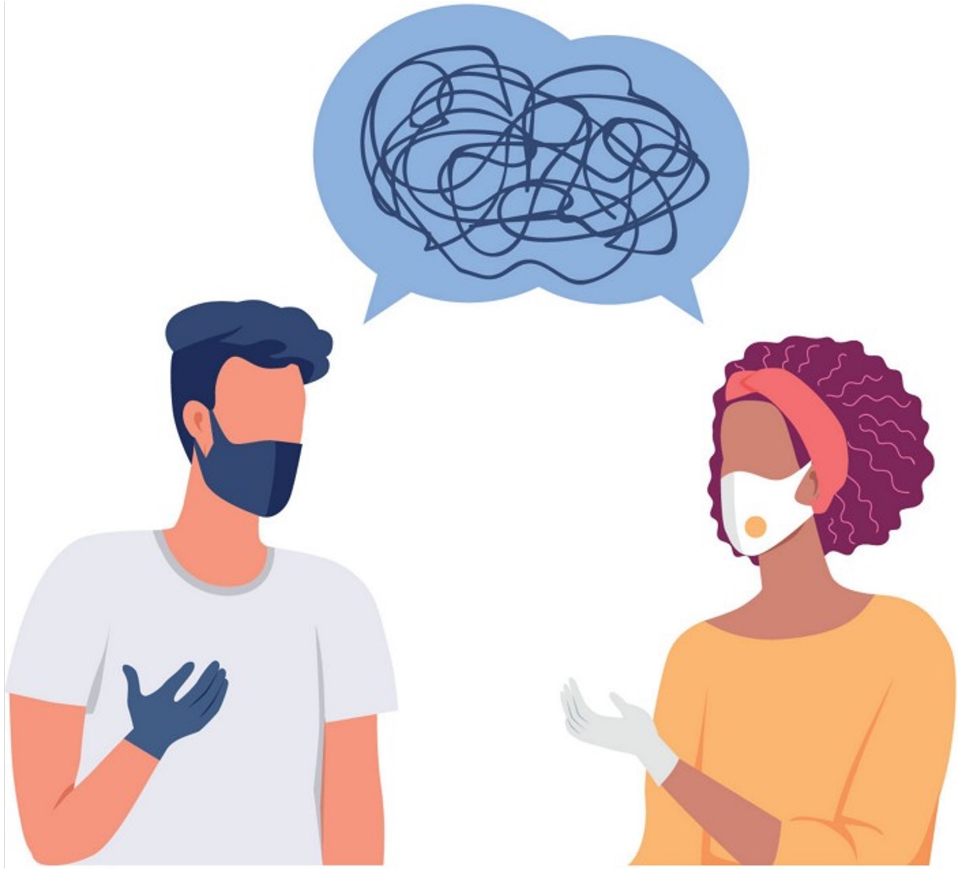

cen.acs.org

### When faced with a difficult circumstance…

- **Identify a safe person** to talk to
  - Friend in lab
  - Friend or family outside of lab
  - Someone in PREP
  - Supervisor
  - Non-supervisor mentor
  - Ombuds
    - an neutral official appointed to handle conflicts against institute

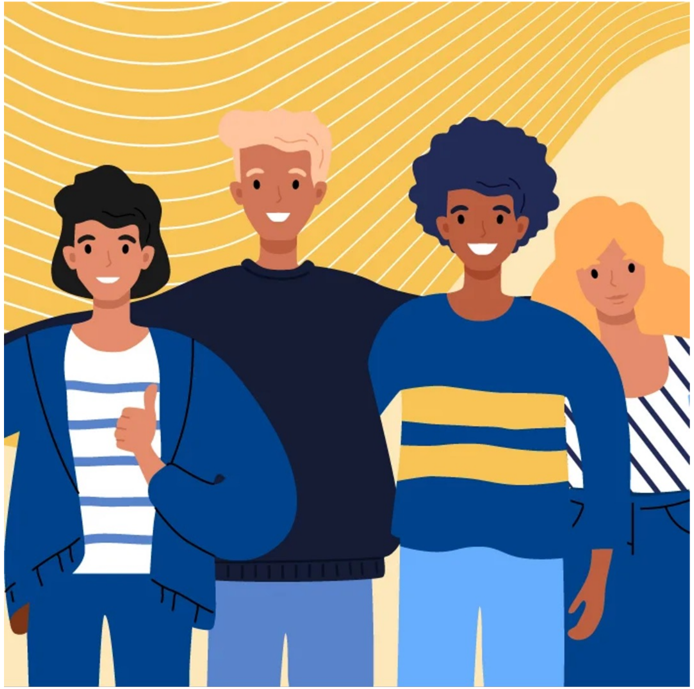

scienceofpeople.com

### Start the conversation

- Determine the **best way for you to communicate** the issue
  - Write it out
  - Talk to a friend/colleague/mentor
- Set up **time to meet**
  - Email
  - Direct conversation
- Share the **reason** for the conversation

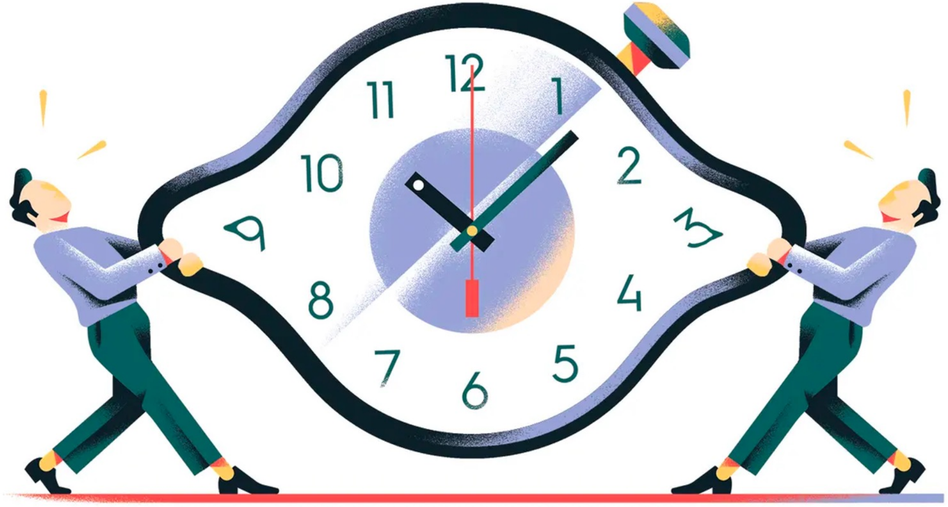

theguardian.com

### Share the reason for the conversation

- Share what happened/your perspective
  - How did it make you feel? And why?
  - **Use "I" statements** about how you feel
    - I felt uncomfortable when "xyz" said he "wouldn't choose me as his girlfriend". I thought it was inappropriate and disrespectful.
  - Share what your goals for the meeting are
  - What actions you **want** and **don't want** as a result

### Have a discussion

- Listen to their perspective
  - Was this a misunderstanding?
  - Is there additional information?
  - Ask questions without being defensive
  - **Clarify goals** of the conversation
- What solutions can you both agree on?
  - What actions can be taken?
  - Agree on **accountability measures**

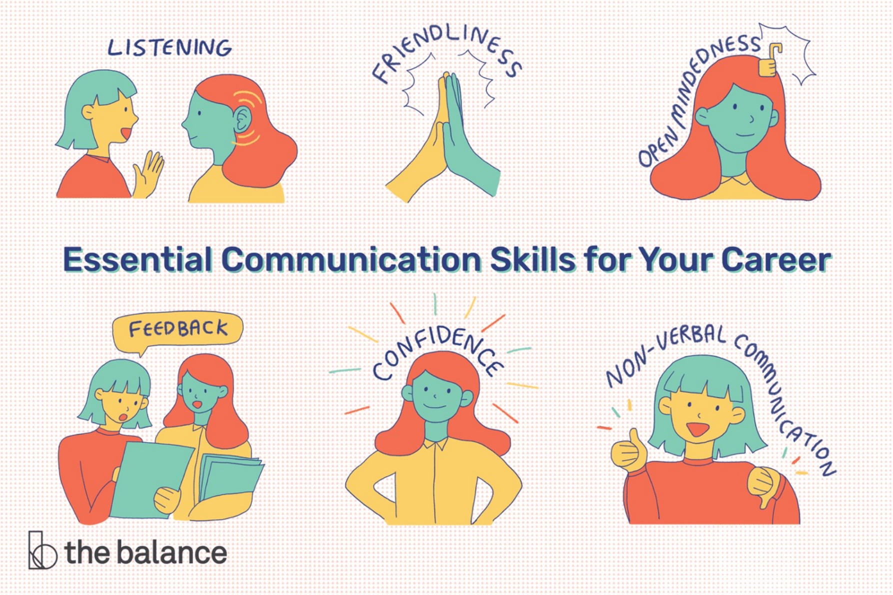


thebalancemoney.com

### Case study #1: Evaluate the situation

1) You notice your lab mate is **claiming your ideas as their own** and taking all the credit for your work

1) You meet with your PI and they make **harsh comments** about your productivity/progress

1) You go to a collaboration meeting and another PI makes a comment about you to the room that is a **microaggression**

### Case study #1: Evaluate the situation

1) You notice your lab mate is **claiming your ideas as their own** and taking all the credit for your work
2) You meet with your PI and they make **harsh comments** about your productivity/progress
3) You go to a collaboration meeting and another PI makes a comment about you to the room that is a **microaggression**

- **How serious are these situations to you?**
  - **Rank them from 1-5**
- **Who would you talk to about the situation?**
- **What steps would you take?**
- **How do power dynamics affect each situation?**

### Case study #2: Think of a time you’ve been in a difficult situation

- How serious was this situation to you?
  - Rank it from 1-5
- Who did you talk to or who would you have wanted to talk to?

### Case study #2: Think of a time you've been in a difficult situation

- How serious was this situation to you?
  - Rank it from 1-5
- Who did you talk to or who would you have wanted to talk to?

- Pair up with a peer mentor and develop some **"I" statements** to practice "sharing the reason for the conversation"

### Resources

- List appropriate institutional resources such as the office of ombuds, office of institutional equity, title IX office, graduate programs, etc

- How to have difficult conversations in the workplace
  - https://www.blackenterprise.com/handle-difficult-conversations-at-work/
  - https://cen.acs.org/careers/career-tips/difficult-conversations-work/99/i4
