## Supplemental Data 3 for "Deepening biomedical research training: Community-Building Wellness Workshops for Post-Baccalaureate Research Education Program (PREP) Trainees"

### COURAGEOUS CONVERSATIONS DISCUSSION PLANNER

#### INITIATE the Conversation

- Use a neutral phrase to open and be mindful of your body language.
- *"I'd like to set up a time to talk with you about our working relationship. Is there a time in the next week or so that would work for you?"*
- [Draft your planned comments here]:

#### SHARE the Purpose of the Conversation

- **Share your intention**. Explain what you do want, and what you don't want from the conversation.
  - *"I am concerned about how things have been going and want to be proactive in seeking to get things back on track. I am hoping this can be a constructive conversation that doesn't leave anyone feeling blamed."*
- **Share your observation and its impact.**
  - Use "I" statements about how you feel and avoid blaming others.
  - "It's important to me to discuss this proactively because I value our shared work and want us to work well together."
  - *"There have been a couple of recent instances where it was clear I did not meet your expectations, and which left me feeling misunderstood and confused."*
  - *"For instance, when _____ happened, I felt _____ because __________. This was concerning to me because I don't feel it reflects my professionalism or commitment to this work. I am confident I can be effective in meeting expectations but need to know clearly what they are."*
  - [Draft your planned comments here]

#### SEEK to Understand

- **Get curious. Gather new information.**
  - *"I'd like to understand this from your perspective. Could you share your own perspective of what happened?"*
  - [Draft your planned comments here]
- **Listen.** Listen genuinely and with an openness to hearing feedback.
  - Ask clarifying, but not defensive, questions.
  - Promote more discussion, i.e. "Could you say more about that?"

#### DEVELOP Solutions

- **Clarify Expectations and Outcome**
  - *"My goal today is for us to get back to a place where we all feel we're working well together. I'm hoping we can use this conversation to make a plan for how we can do that."*
  - [Draft your planned comments here]
- **Inquire About Ideas, Listen and be open.**
  - *"Do you have any specific suggestions that you feel would help?"*
  - *"That's interesting, thank you for suggesting that. Do you have any other thoughts?"*
- **Share any ideas you may have.**
  - *"I had some ideas as well. Would you be open to talking them through together?"*

#### AGREE on Solutions and Next Steps.

- **State your understanding of any agreements reached.**
  - *"As I understand it, we've agreed that we'll ________________."*
- **Check their understanding of the agreement.**
  - *"Does that match your understanding of what we've discussed?"*
- **Determine measurement and accountability.**
  - *"Can we set a time to check in and ensure that this is working as planned?"*

#### CLOSE with Respect.

- **Summarize agreements and next steps.**
  - *"I will send an email that attempts to capture what we've come up with today."*
- **Share your confidence in what you've accomplished together.**
  - *"I feel good about what we've agreed to today and am confident that this will be effective in helping us work better together."*
- **Say "Thank you."**
  - *"Thank you for taking the time to connect about this today. I appreciate and value you as supervisors/colleagues."*
