## Supplemental Data 4 for "Deepening biomedical research training: Community-Building Wellness Workshops for Post-Baccalaureate Research Education Program (PREP) Trainees"

### Courageous Conversations Reflection Worksheet

#### What are Our Stories?

| **My Story**<br>What is the problem/issue from my point of view? | **Their Story**<br>What is the problem from their point of view? |
|---|---|

#### Our Contributions

| **My Contribution**<br>How have I contributed to the current situation? | **Their Contribution**<br>How have they contributed to the current situation? |
|---|---|
| **Impact**<br>What impact might this situation have had on them? | **Impact**<br>What impact has this situation had on me? |

Adapted from Triad Consulting 2009, Difficult Conversations www.diffcon.com

#### Feelings at Play?

| **My Feelings**<br>How do I feel about the situation and why? | **Their Feelings**<br>What might they be feeling? Why? |
| --- | --- |

#### Identity

| **My Self Image**<br>What do I fear this situation says about me? | **Their Self Image**<br>What might the situation say about them that would be upsetting to them? |
| --- | --- |

| **What I <u>don't</u> want from this conversation** |
| --- |
| **What I <u>do</u> want from this conversation** |
