## Supplementary material for "Deepening biomedical research training: Community-Building Wellness Workshops for Post-Baccalaureate Research Education Program (PREP) Trainees": S12 Slidedeck for Time Management

### Overview

- This workshop is centered around being cognizant of and improving our time-management skills. We will focus on common pitfalls and strategies to improve our skills and finish with one on one peer coaching to make an action plan moving forward.

### Pre Work

- Reflect on your current time management strategies and struggles. Specifically, think back to this past week, were you able to get done what you needed? What time-management strategies did you use? How did you feel throughout the work week and how are you feeling for this upcoming week?

### Post Work

- For post work, we would like everyone to meet with a Peer Mentor and discuss (1) if you've implemented any new time management strategies, (2) how you are feeling throughout the previous work week, (3) how you are feeling for the upcoming weeks in regards to time management.

### Time Management

Johns Hopkins PREP Workshops

### Overview of the day

- Reflection
- Time management struggles/pitfalls
- Useful Techniques & Tools
- Self-Care Techniques
- Peer Coaching

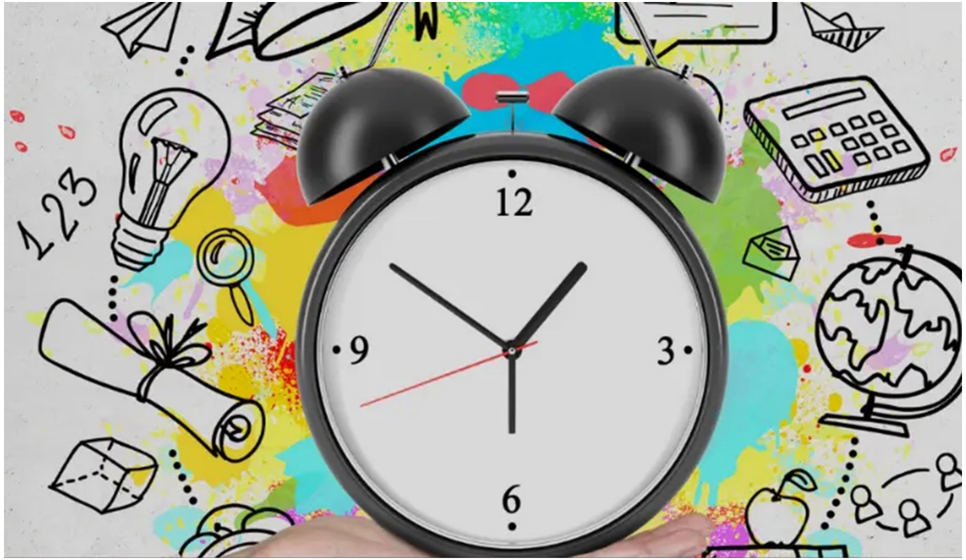


https://www.adda247.com/school/time-management-tips-for-student/

### Group Reflection

**Bonita**: Work-life balance, Saying No

**Vincent**: Overestimating how much I can accomplish

Small Group Discussion: What do you struggle with?

### Common Time Management Struggles

- Being a perfectionist
- Lack of vision
- **Procrastination**
  - **~20% of adults are chronic procrastinators[1]**
- **Stress**/Fear of failure
- Not being able to concentrate/focus
- Being unmotivated

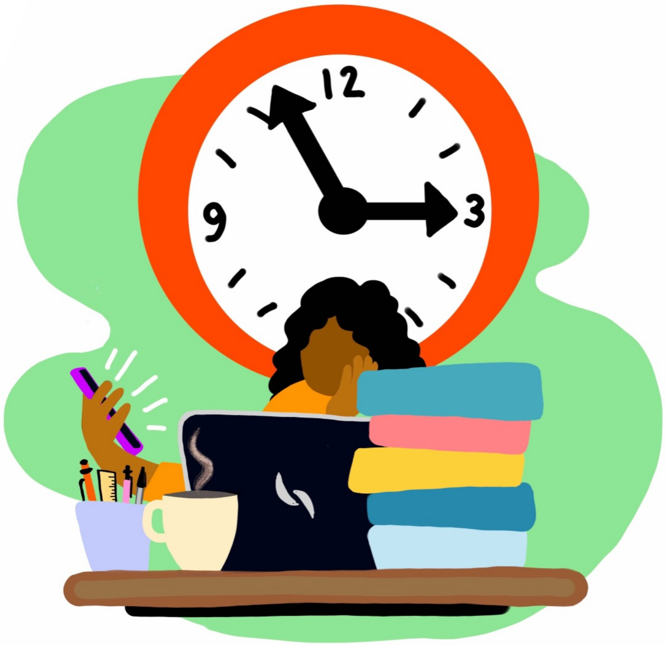


https://dailytrojan.com/2021/11/30/stop-procrastinating-tomorrow/

[1] Gustavon *Psychol. Sci,* 2014

### Tips/Strategies

- **Make an action plan**
- Control your environment
- Set boundaries
- Work towards your strengths
- **Eat the frog**
- Stay organized
- Ask for help/delegate

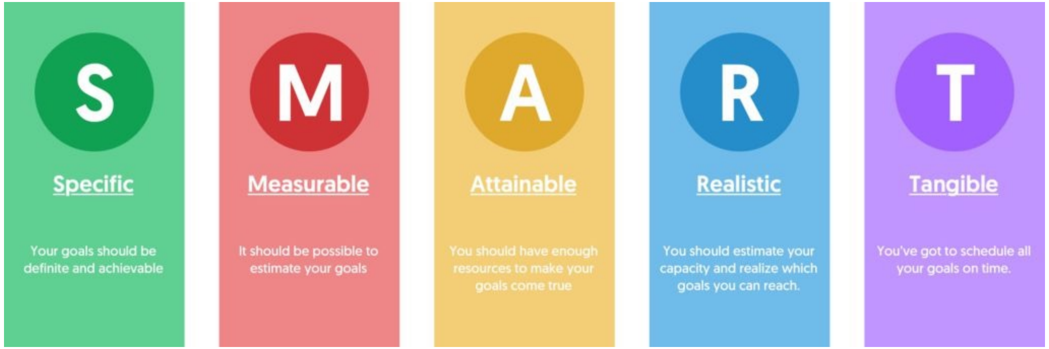


https://everhour.com/blog/time-management-strategies/

### Make an Action Plan

- An action plan outlines the steps to achieve a specific goal.
- A tool to monitor and track your progress.
- **Tools to develop an action plan.**
    1. Flowcharts
    2. **Gantt chart**
    3. Mind mapping
    4. Spreadsheets
- **What to include?**
    1. Expected outcomes
    2. List of tasks
    3. Required resources
    4. Timeline

**Gantt chart**

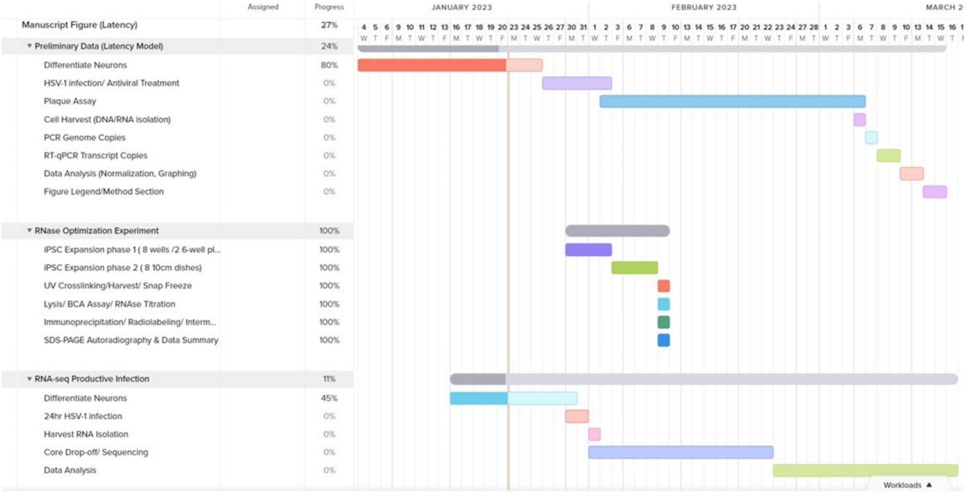

### Eat the Frog

- Phrase coined by Mark Twain, "…if you have to eat a live frog, do it first thing in the morning and nothing worse will happen to you for the rest of the day."
- Identify a challenging task for the day, and do it in the morning (**most productive time of the day**)
- Don't plan the task too far in advance!
- Try to make this a habit!

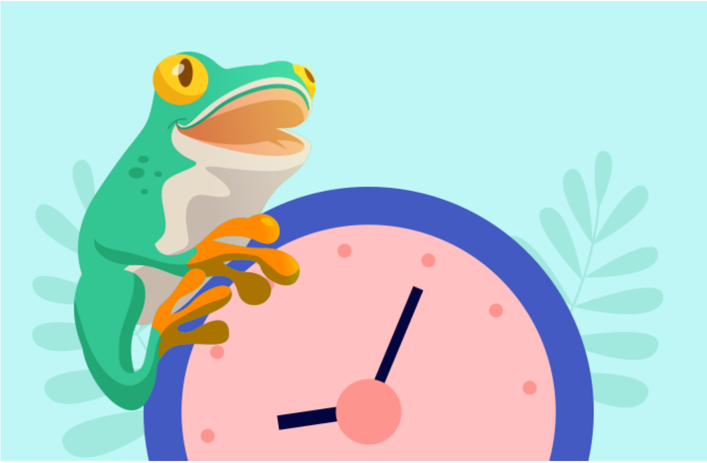

https://www.teamly.com/blog/eat-that-frog-time-management/

### Tips/Strategies

- **Pomodoro Technique**
- Take breaks
- Be careful with Multi-tasking
- **Break down goal into smaller tasks**
- Remember your "Why"
- Reward yourself!

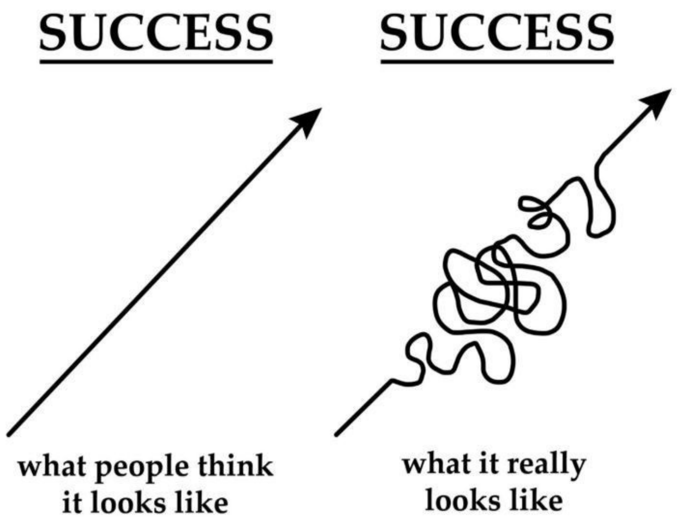


https://beyouonlybettereveryday.medium.com/the-power-of-taking-small-steps-to-achieve-your-goals-3b56d58caf9c

### Pomodoro Technique

- Get a to-do list and a timer
- Set your timer for 25 minutes, focus on a single task until the timer rings
- Take a 5-minute break!
- Repeat 3 times and then take a longer 30-minute break

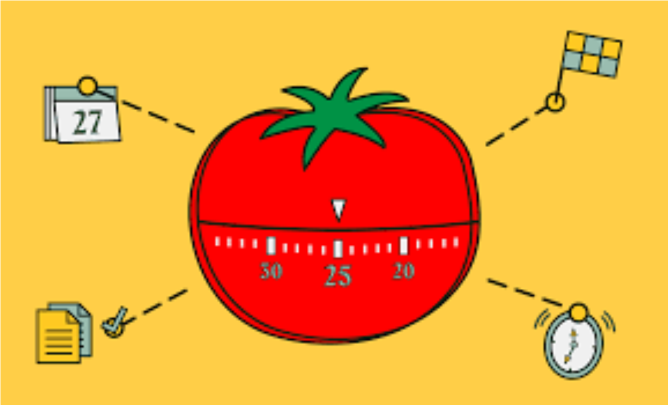


https://www.lockcard.app/post/why-the-pomodoro-technique-is-effective-for-studying-and-productivity

### Break Down Goal into Smaller Tasks

- **Breaking down a goal helps to make the goal more manageable and achievable.**
  1. Can visualize steps.
  2. Can prioritize easier.
  3. Track progress.
  4. Can adjust plan.
  5. Feel less overwhelmed.

- **How to breakdown a goal**
  1. Ask “What” and “How”
  2. Determine main stages.
  3. What needs to be completed first?
  4. Look at each stage, divide into smaller tasks.
  5. Create a timeline.

Group Discussion: What techniques work for you? Which do you want to try?

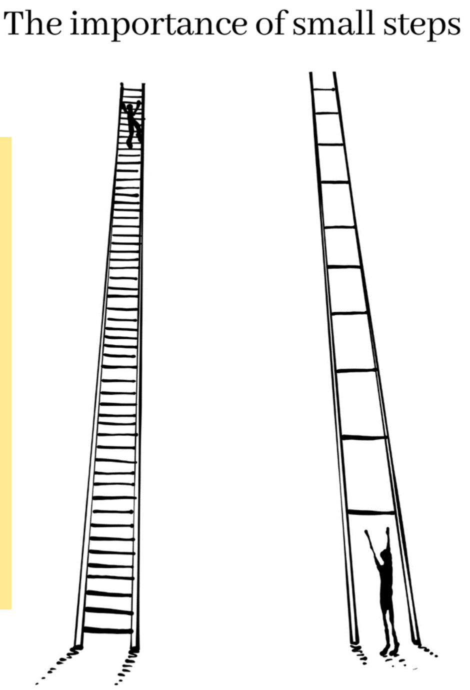


https://www.psychedconsult.com/the-importance-of-small-steps/

### Prioritizing Ourselves!

- Self-care/ Giving yourself breaks
- Saying No
- Work/Life balance
- Asking for help

Group Discussion:
How do you practice self-care?

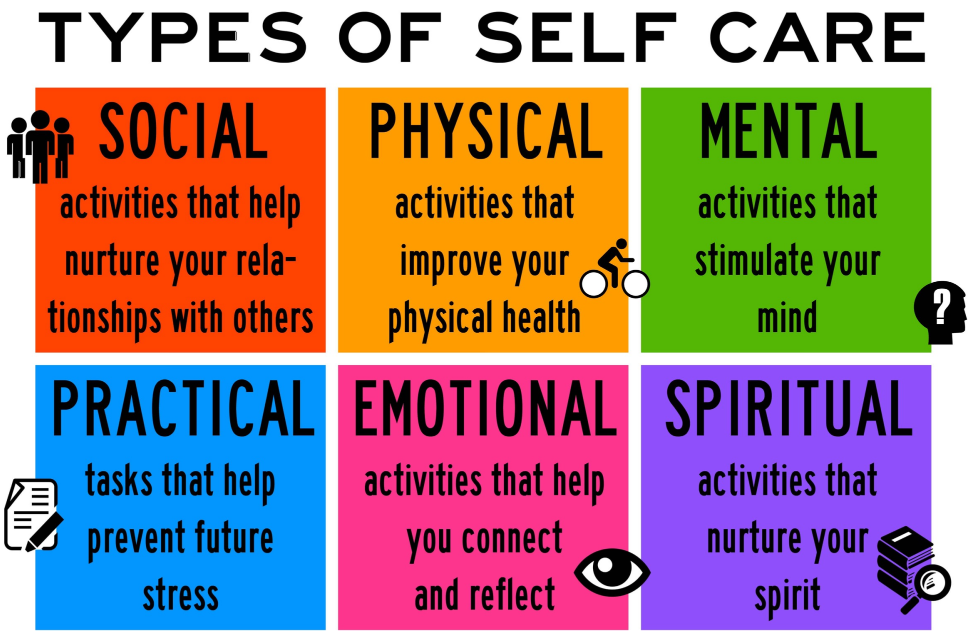


https://reachoutrecovery.com/what-is-self-care-today/

### Time Management in a Crisis

- Stress can impact how much & what type of work we can perform
- **Types of Stressors**:
  1. Upcoming Exam ( i.e. MCAT)
  2. Graduate School Interviews
  3. Unexpected Event
  4. Family/Personal/Health
  5. Research Progress
- **Symptoms of Stress**:
  1. Fatigue
  2. Mental Block
  3. Trouble sleeping
  4. Memory lapse
  5. Poor focus
- **What to Do:**
  1. Recognize stress symptoms.
  2. Take a break/Self-care
  3. Focus on stress management
  4. Modify action plan/**do what you can**

Group Discussion: What techniques work for you during a crisis?

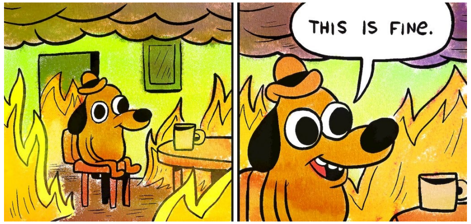


https://www.nytimes.com/2016/08/06/arts/this-is-fine-meme-dog-fire.html

### Peer Coaching

- What do you struggle with in terms of time management?
- What would you like to see or experience moving forward?
- What have you tried so far?
- What has worked for you in the past?
- What support do you want to move forward?

**Stop Start More Less Grid**

| | |
|---|---|
| What will you STOP doing? | What will you START doing? |
| What will you do MORE of? | What will you do LESS of? |

### Additional Resources

- Guidelines to Draw a timeline of your PhD
  - https://academiac.net/2018/11/20/guidelines-to-draw-a-timeline-of-your-phd/
- Google Calendar
  - https://calendar.google.com/
- Pomodoro Timer
  - http://www.tomatotimers.com/
- Team Gantt
  - https://www.teamgantt.com/
- ADHD Jessie @adhdjesse
  - https://www.extrafocus.net/
- Good Notes
  - https://www.goodnotes.com/

**Eisenhower Matrix**

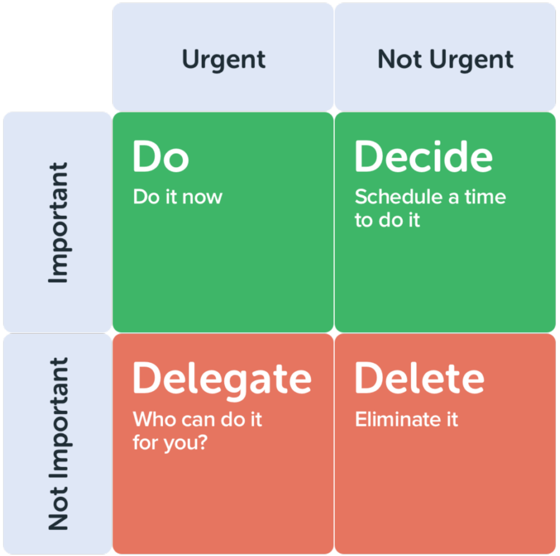


https://www.tameday.com/time-management-strategies/
