## Supplementary material for "Deepening biomedical research training: Community-Building Wellness Workshops for Post-Baccalaureate Research Education Program (PREP) Trainees": S13 Slidedeck for Owning your Identity

### Overview

The goal for this Owning Your Identity Wellness Workshop is to encourage PREP Scholars to reflect on whether the professional opportunities they are pursuing align with their needs to feel whole, valued, and happy. This also involved understanding that life circumstances, values, and obligations change with time.

### Prework

- **Please reflect on your finalized on ongoing journeys of applying to Ph.D./MSTPs. With that reflection, think about how your identities/values have shaped your considerations for graduate programs (This includes all aspects: program structure, cost of living, recreational activities available, access to nature, access to a big city, potential for discrimination, proximity to family/friends/partners, etc.) If comfortable, please be ready to discuss your reflection as a group.**

### Postwork

- Visualizing what your dream life is, design a slide with words and images that reflect your future self in the next 5 years and 10 years. How do you anticipate your identity/values to change on this journey?

### Owning your identity

Johns Hopkins PREP Workshops

#### What immediately comes to mind when you think of the factors that make up your identity?

### Share your Pre-Work!

### Who or what defines you?

***"Definitions belong to the definers, not the defined."***

***(Toni Morrison, "Beloved," 1987)***

*"How do I know who I am or where I am? How could a single wave locate itself in an ocean?"*
*- Rumi*

### What is Identity?

- Definition: a person's sense of self, established by their qualities, beliefs, personality traits, appearance, and expressions
  - It's how you define yourself and, also how others define you

### How can we define who we are?

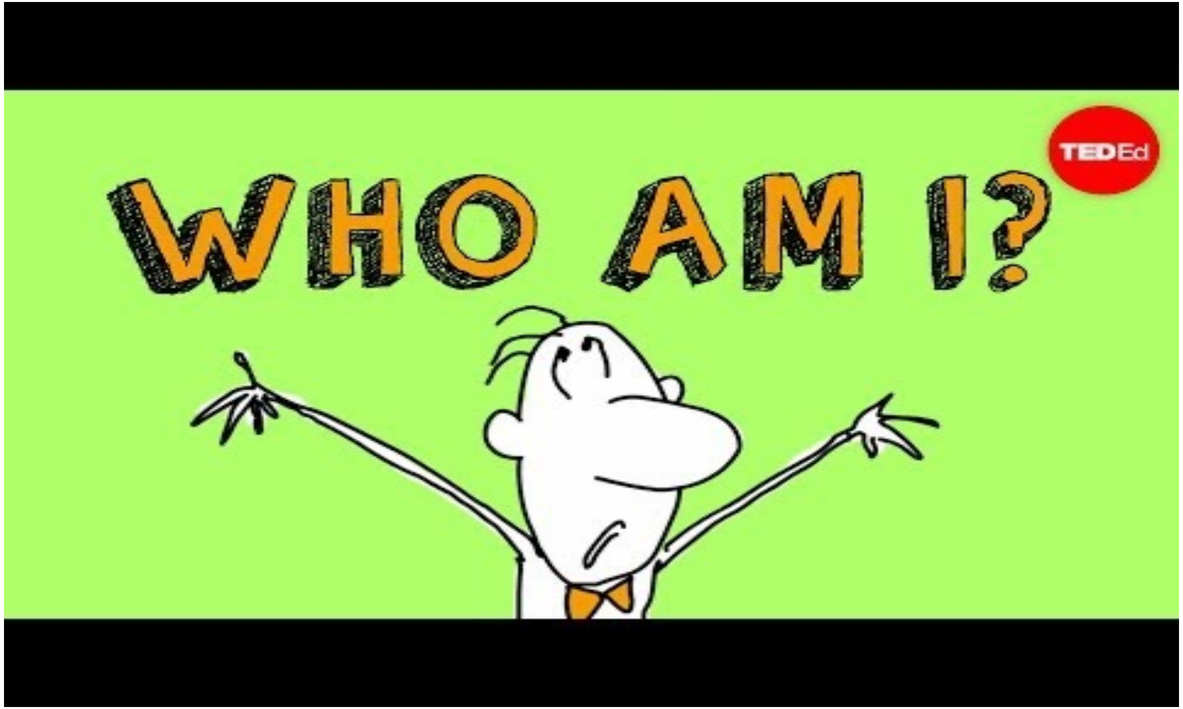


Link: https://youtu.be/UHwVyplU3Pg

### Identity is fluid

- You have ***grown*** from the person that you were yesterday and ***will be a different person tomorrow*** than you are today
  - Ex: You mature, values and priorities change, boundaries to protect time, mental health, and establish healthy self-advocacy shift
- Just because ***you change*** does not mean that you are inauthentic

### Who have you *been*?

- Imagine your middle school self: who were they?
  - Draw you’re an image of yourself in middle school. Include icons and words that represent your ideals and values

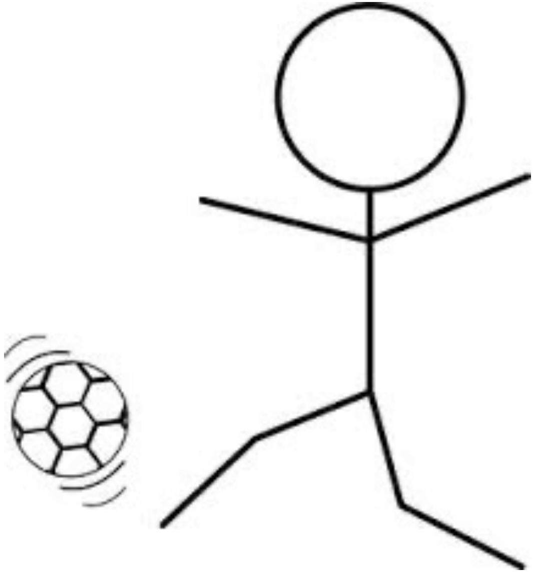

### Who are you *now*?

- Imagine yourself in the past year or two: who are you?
  - Draw you're an image of your current self. Include more icons and words that represent your ideals and values

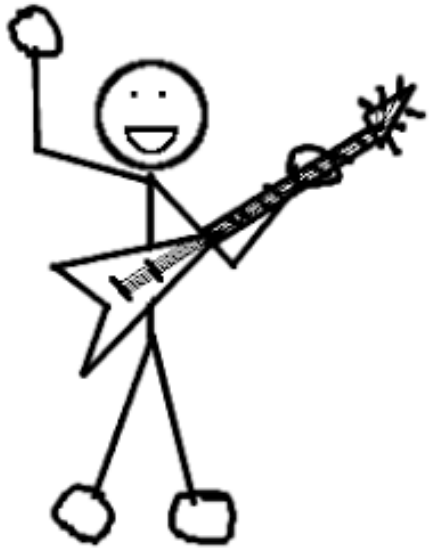

### Who will you *become*?

- ❑ Imagine the possibilities for your future self in the next 5 years
  - ❑ Draw you're an image of your future self. Include more icons and words that represent your ideals and values

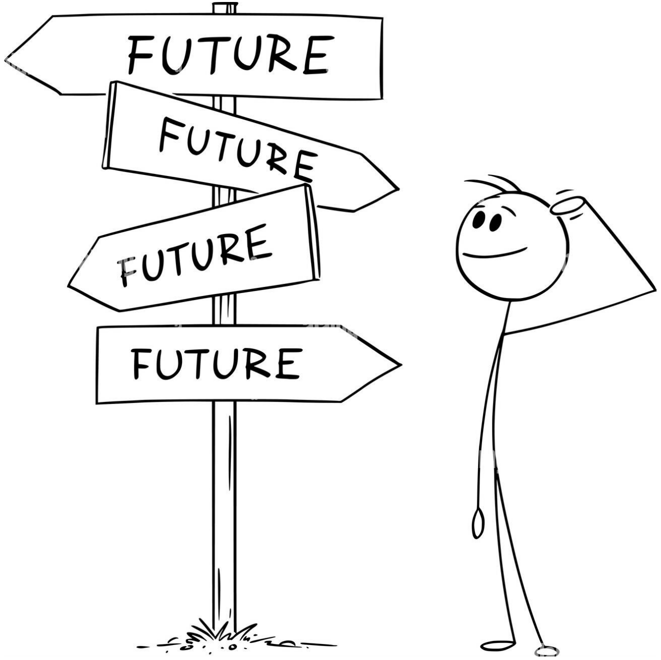

### Owning your identity is the center of well-being

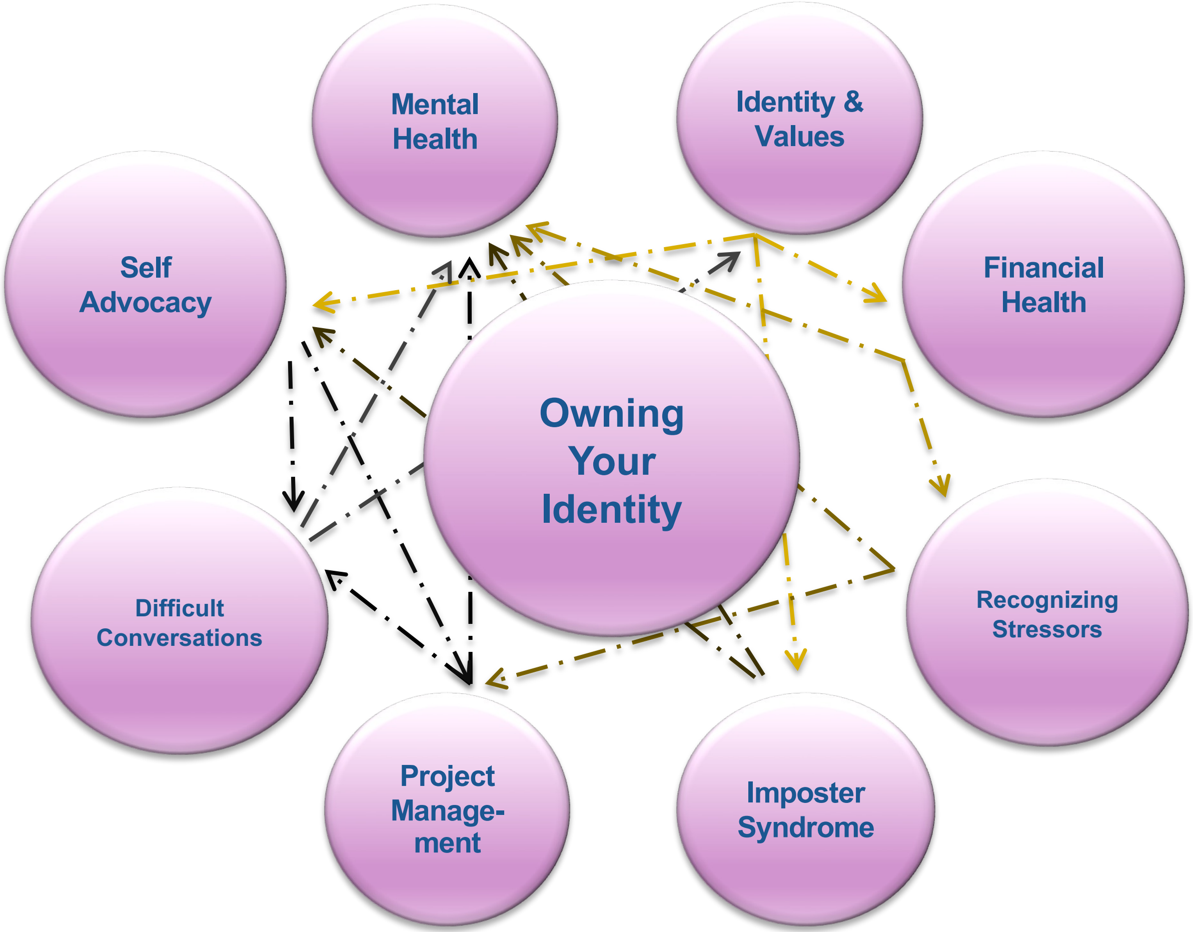

### Creating the Space that Aligns with You

- ❑ Self-reflection of what you prioritize based on all the external and internal factors that comprise your identity (5 mins)
  - ❑ What priorities remain constant?
  - ❑ What is flexible?
- ❑ Share your self-assessment with others

### What feeds you?

- **We receive and give energy** to go about **tasks and throughout interactions** in our daily lives.
- *Tasks and interactions* with others, whether active or stationary, *can be emotionally and physically invigorating or draining based on how they **align** with you*
    - What are some energy givers in your life? (Things that bring joy that connect with your identity)
    - What are some energy takers in your life?
- Share your self-assessment with others
- Reflection helps you strengthen your sense of identity

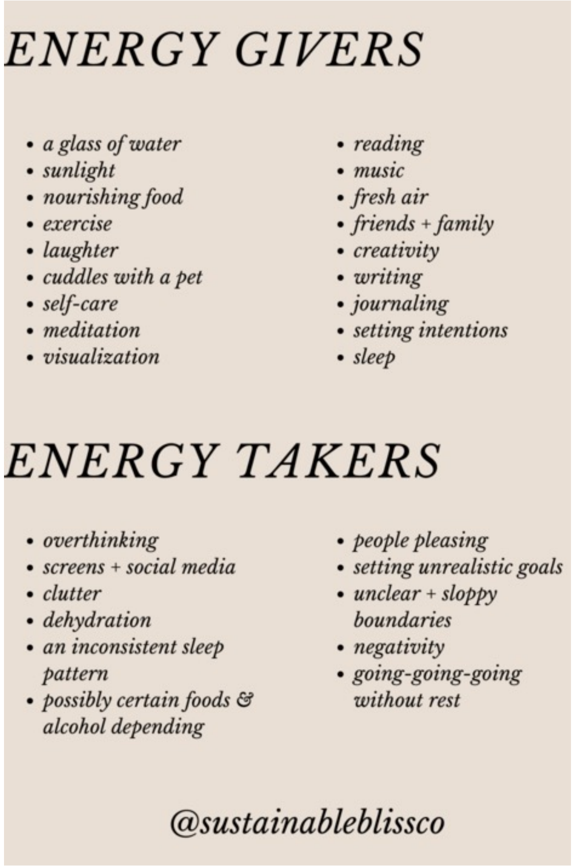

### Case Study: Owning your identity is the center of well-being

Nisi grew up in a farming town. Their parents are famers, and their grandparents were farmers. From a very young age, Nisi handled horses and cows. In whatever spare time Nisi could find, they participated in volunteer work in town. Most elders saw Nisi as a quiet, well-mannered young person with exceptional farming skills  and a giving heart. Growing up, Nisi also enjoyed playing in a rock band with their friends and enjoyed being a leader. After high school, Nisi moved away to pursue a career in emergency medicine and perform with their band in larger cities.

**From what you have learned about Nisi, are their choices in career, location, and recreational activities in line with their identity? Why or why not?**

- **What may others think of their choices?**
  - **Does this matter?**
- **Reflect on your own journey.**

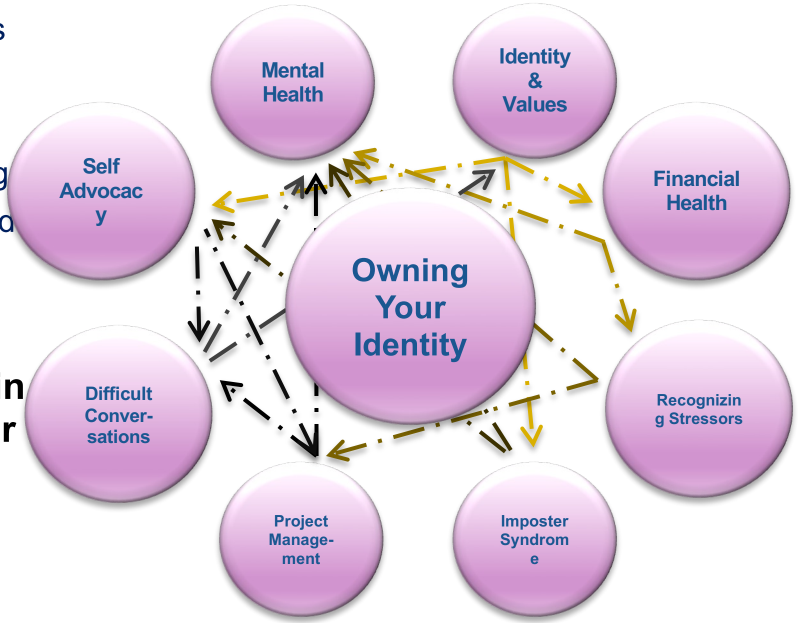

### Who or what defines you? (revisited)

***"Definitions belong to the definers,***
***not the defined."***

***(Toni Morrison, "Beloved," 1987)***

### Resources

**You will *continue* to change! Reflecting and reassessing can help you gain clarity of your current needs. Use the resources that bring you energy and peace. These can include the following:**

- **Mental Health Professionals**
- **Friends and Family/Partners**
- **Primary Literature and books**
- **Things that bring you joy and that connect with your identity.**
- **Identifying mentors and colleagues with shared identities/ those that fully see and value your full identity and humanity.**

### List of Resources

- **Rachel Crowell *Bringing the Whole Self to Science.* Symmetry (2021)**
  – https://www.symmetrymagazine.org/article/bringing-the-whole-self-to-science
- **Nedra Glover Twabb *Set Boundaries, Find Peace: A Guide to Reclaiming Yourself (2021)***
  – *https://www.nedratawwab.com/set-boundaries-find-peace-1*
- **Najwa Zebian *Welcome Home (2021)***
  – https://www.welcomehomebook.com
- **Guitiérrez y Muhs, Flores Niemann, González, and Harris *Presumed Incompetent: The Intersections of Race and Class for Women in Academia***
  – https://www.amazon.com/dp/0874219221/?coliid=I27H22WAAYTBNJ&colid=9QLV9E3LYY4K&psc=1&ref_=list_c_wl_lv_ov_lig_dp_it
- **Kerry Ann Rockquemore *The Black Academic's Guide to Winning Tenure- Without Losing Your Soul* (2008)**
  – https://www.amazon.com/Black-Academics-Winning-Tenure-Without-Losing/dp/1588265889/ref=pd_bxgy_vft_none_img_sccl_2/131-1825531-4220366?pd_rd_w=D1tYN&content-id=amzn1.sym.26a5c67f-1a30-486b-bb90-b523ad38d5a0&pf_rd_p=26a5c67f-1a30-486b-bb90-b523ad38d5a0&pf_rd_r=XARRZ4JEEHE13TPHDBKH&pd_rd_wg=N33i0&pd_rd_r=eb7f12a2-0dc9-412e-86de-8f55ecfd2cb6&pd_rd_i=1588265889&psc=1
