## Supplementary material for "Deepening biomedical research training: Community-Building Wellness Workshops for Post-Baccalaureate Research Education Program (PREP) Trainees": S14 Scholar Exit Survey Questions

**Scholar Exit Survey Questions Only**

*LQ = Likert question; For likert questions (LQ): 1 = Bottom of Scale; 5 = Top of Scale*

*FR = Free Response*

1. Select the top three MOST impactful workshops for you? (pull-down menu)

2. Select the three LEAST impactful workshops for you? (pull-down menu)

3. By the end of the year, how comfortable were you sharing during the Workshops? (LQ)

4. Describe any factors that increased your comfort when sharing during the workshops. (FR)

5. What can we do better next year to promote more open dialog during the workshops? (FR)

6. Did you ever reach out to a Peer Mentor this year for help or advice? (Yes/No)

7. How would you rank your comfort level asking for help from the Peer Mentors? (LQ)

8. What factors increased your comfort level reaching out to Peer Mentors? (FR)

9. What could we improve on next year to increase your comfort interacting with Peer Mentors? (FR)

10. Have you put into practice any topics covered in the Workshops? (Yes/No)

11. Explain what Workshop skills you have put into practice. (FR)

12. Have the workshops spurred you to bring up any topics in either your 1:1 meetings with Kathy, your PI, or in-lab mentor? (Yes/No)

13. Have you changed how you interact with your lab mates or mentors changed because of what is covered in the workshops? (Yes/No)

14. Have you discussed the Workshop topics with our PREP scholars outside of the workshops? (Yes/No)

15. Have your criteria for choosing future mentors changed as a result of the workshops? (Yes/No)

16. How likely are you to seek out a community similar to the Peer Mentors in your next step? (LQ)

17. How much of a priority was attending the workshops to you when setting up your schedule? (LQ)

18. Identity and Values Workshop. Do you feel that the Values and Identity Workshop has helped you consider whether your values are reflected as you decide the next step of your career? (LQ)

19. Identity and Values Workshop. Since the Values and Identity Workshop, have you re-evaluated your career goals based on your personal values? (Yes/No)

20. Financial Health Workshop: Do you feel that the presentation about finances provided you with any new information or a framework from which you could do your own research on how to handle your money? (LQ)

21. Financial Health: Since the workshop, are you more proactive in handling your finances? (Yes/No)

22. Recognizing Stressors: Since the workshop, do you feel better equipped at recognizing signs of emotional and physical stress manifestations in yourself? (LQ)

23. Recognizing Stressors: Have you been able to implement/reinforce healthy short-term and long-term stress management tools? If not, what changes to the workshop would help make identifying stressors and developing healthy coping mechanisms more effective? (FR)

24. Imposter Syndrome: Were the discussions held around imposter syndrome and the mislabeling of discrimination in science as imposter syndrome useful? (Yes/No)

25. Imposter Syndrome: Did you find the Peer Coaching work useful? What did you like or did not enjoy about it? Do you think it is most useful to do during NPM workshops? (FR)

26. Time Management: Did this workshop help you improve your time management skills? (LQ)

27. Time Management: Have you employed any of the time management techniques discussed? If so, what worked? If not, what strategies would you like to try? (FR)

28. Difficult Conversations: Did the difficult conversations workshop equip you with new strategies to have difficult conversations in a professional setting? (LQ)

29. Difficult Conversations: What was most helpful about this workshop? How could this workshop be improved? (FR)

30. Self Advocacy: Since the self-advocacy workshop, how often would you say you use assertive language in your lab? (LQ)

31. Self Advocacy: Has this workshop helped identify your boundaries for assertiveness? (Yes/No)

32. Mental Health: Since the mental health workshop, do you feel more comfortable seeking professional help if needed. (LQ)

33. Mental Health: How has this workshop changed your perspective on mental health? (FR)

34. Owning your identity: Do you feel like the workshop equipped you with new tools or a new perspective on identity? (LQ)

35. Owning your identity: Since this workshop, have you thought more about how future career plans align or misalign with your identity? If so, in what ways did the workshop impact thinking about your plans? (FR)

36. Last question -- is there anything else you would like to share about the Workshops and the Peer Mentors? (FR)
