## Supplementary material for "Deepening biomedical research training: Community-Building Wellness Workshops for Post-Baccalaureate Research Education Program (PREP) Trainees": S15 Peer Mentor Exit Survey Questions

**Peer Mentor Survey Questions**

*LQ = Likert question; For likert questions (LQ): 1 = Bottom of Scale; 5 = Top of Scale*

*FR = Free Response*

1. How equipped did you feel to discuss topics raised by PREP scholars? (LQ)

2. What changes would you suggest to improve Peer Mentor preparation/efficacy? (FR)

3. Did you ever meet with any scholars outside of the PREP official events? (Yes/No)

4. If you did meet with scholars, who initiated the meeting? (FR)

5. Did you feel like you were able to make meaningful connections with any scholar(s)? (FR)

6. What could we do next year to help increase a sense of community between scholars and peer mentors? (FR)

7. How comfortable were you sharing relevant aspects of my personal journey during the workshops? (LQ)

8. What could we improve on to create a safer space to share? (FR)

9. I felt confident in my ability to facilitate my workshop. (LQ)

10. What factors/resources would help improve your facilitating? (FR)

11. I would have benefited from similar workshops earlier in my training. (LQ)

12. I benefited from these workshops at this step in my training. (LQ)

13. I think these workshops would be beneficial to those in graduate-level training programs. (LQ)

14. The wellness workshops have changed my mentoring style. (Yes/No)

15. At earlier steps in my training, I was a participant (aka Scholar) in either a PREP program or similar program. (LQ)

16. Why did you choose to be a PREP mentor? (FR)

17. I would recommend being a Peer Mentor to others.
