## Supplementary material for "Deepening biomedical research training: Community-Building Wellness Workshops for Post-Baccalaureate Research Education Program (PREP) Trainees": S1 Slidedeck for Values & Goals Workshop

### Overview

- As this is the first workshop it served dual roles:
  - To introduce the program, set expectations, and establish this group as a safe space.
  - The "Goals and Values" works aims to a sense of community between all scholars and mentors by discussing our values and goals. This helps to understand the experiences of each of our peers, many of which are similar. Additionally, it is important to understand if our values align with our goals and if not, adjust accordingly.

### Pre-work

- Create a single slide with 3 images that represent values you hold. For each image, include a 1-word value it represents and submit to be presented in the session.
- It is meant to be short! We want to distill down our values into their simplest terms to really understand what is important to us.

### Post-work

- Reflect on these 3 questions throughout the workshops this year. Note how the answers may change throughout the course of the workshop series.
  - How have your values influenced your goals?
  - Do your values fit with the goals you have?
  - What questions do you have to ask yourself to determine whether they fit?

### VALUES & GOALS

Johns Hopkins PREP Workshops

JOHNS HOPKINS
SCHOOL of MEDICINE

### Goals of the Wellness Workshops

- Develop life-long skills and practices in the themes of introspection, mental health, community, emotional intelligence, and financial fitness

### HOW

- Introspection to create an individual framework
- Practice "hard" skills and move beyond our comfort zone
- Adapt a growth mindset
- Work together as a community

### Expectations for our community

- Be present
- Contribute
- Respectful

Suggestion – gather input from the group on what showing respect means to them.

### Values Slides

3 photos, each representing a different value

1 word for each photo that reflects your values

Include slides gathered from the group

### Identity deeply influences our values

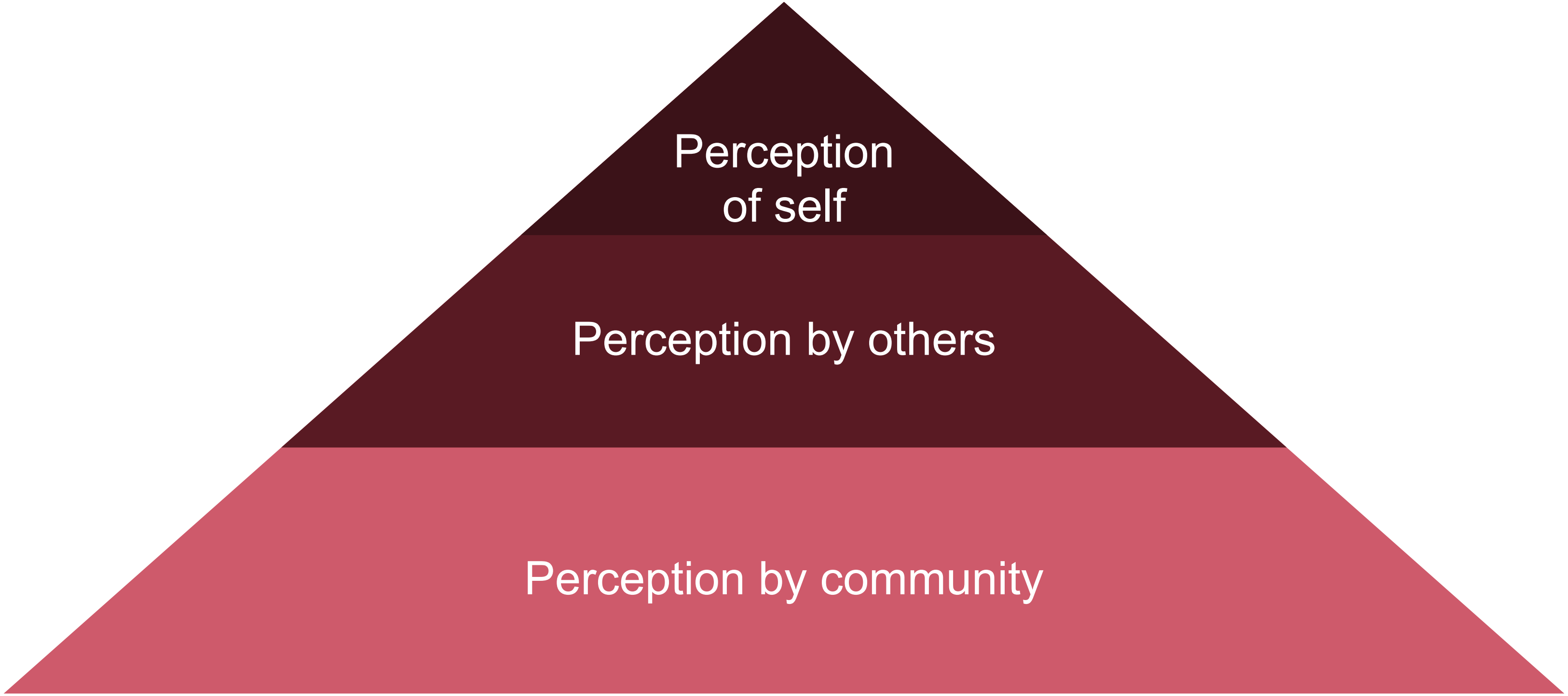

### Identity and values (breakouts/small groups)

- How does your identity impact the way you see yourself?
- How does your identity impact the way others see you?
- How does your identity impact what it means to be in community with others?

### Goal-setting

"Be" is what type of characteristics you want to develop in yourself.

"Do" is what type of things you want to try or do in your life.

"Have" are the material (or non-material) possessions you wish to have.

### Goals (Breakouts/small groups)

- Who do you want to be?
- What do you want to do?
- What do you want to have?

### Reflection Questions

How have your values influenced your goals?

Do your values fit with the goals you have?

What questions do you have to ask yourself to determine whether they fit?

### Overview

- The goal of this workshop is to increase Scholar long-term financial literacy by providing concrete resources and discussion from which they can start/continue their own research into their individual financial situation (slides should be shared as they contain links)

- We do not want to tell them how to spend their money; we want them to reflect on how they currently handle their money and if they will do anything different after this discussion

### Pre-work: speed budgeting exercise

**Building a basic budget relies on 3 things:**

1. Knowing your net income (from all sources)
2. Creating a list of your monthly expenses
3. Calculating a monthly amount of income left over (your "cash flow")

https://www.huntington.com/learn/budgeting/how-to-make-a-budget

**Use the provided worksheet and fill out the following info in ONLY 3 minutes, then be prepared to discuss the day of the workshop:**

- Create a list of your monthly expenses (specific $ amount NOT needed), identify which are fixed vs. variable
  - Fixed: same amount each month
  - Variable: amount can fluctuate each month
- Characterize which are "want" vs. "need" expenses (waying your priorities)
  - Want: trip to Bali
  - Need: groceries

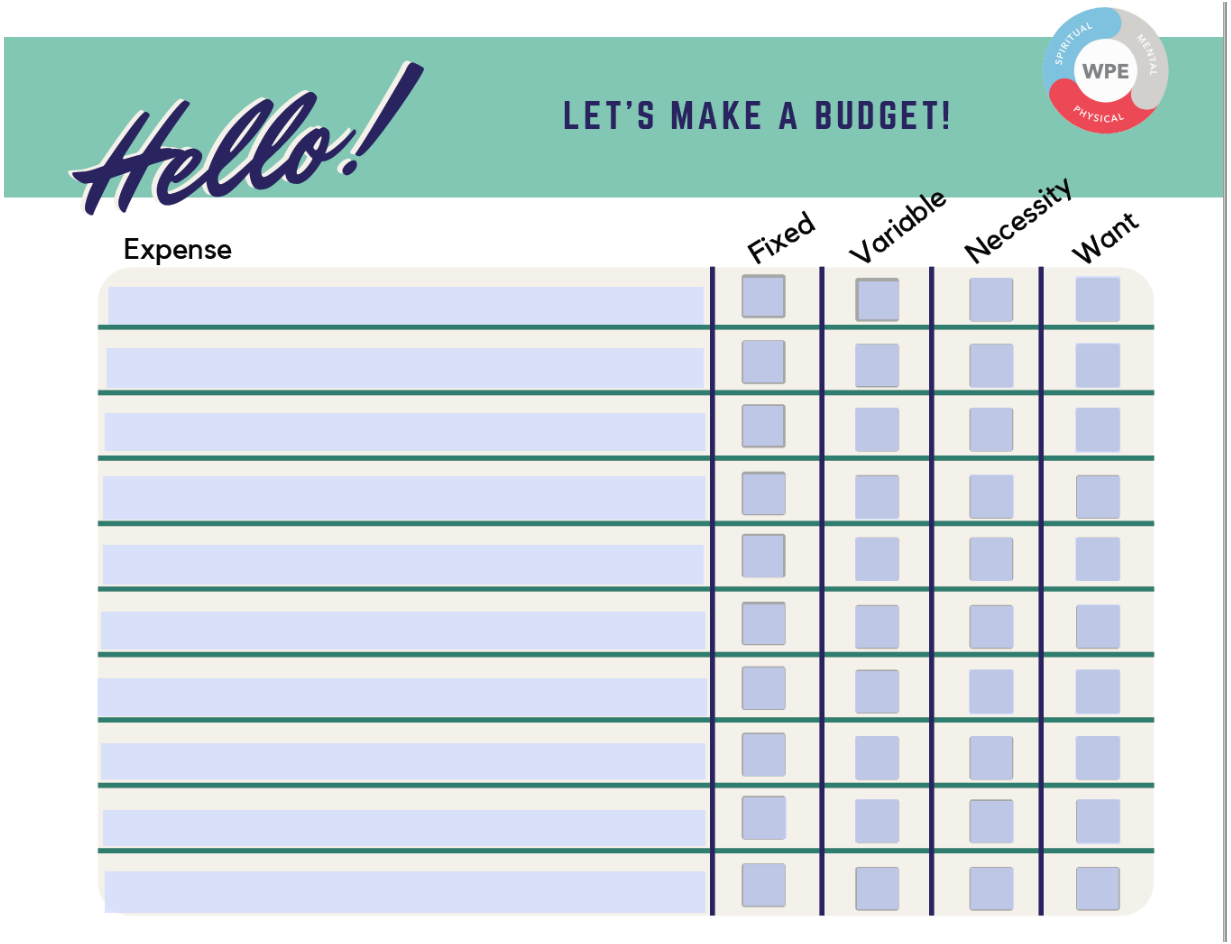


Source: California State University Channel Islands

### Post-work: thinking about credit

**Research the following questions, then discuss with either another PREP Scholar or Peer Mentor:**

- What is credit?
- What is a credit score? And how can you find yours?
- How do you build credit?
- What positively and negatively impacts credit?
- What can credit do for you or allow you to do?

https://studentaid.gov/

Federal Student Aid
An OFFICE of the U.S. DEPARTMENT of EDUCATION

### Future goals and finances :

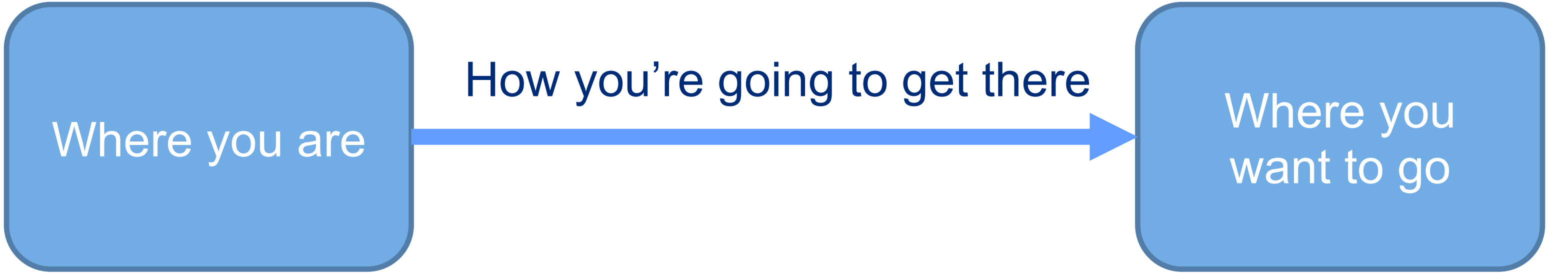


- Do your current finances align with your goals/values?

- What can you do in your current position to enhance your financial wellbeing?
